# Cation-controlled assembly, activity, and organisation of biomimetic DNA receptors in synthetic cell membranes

**DOI:** 10.1101/2025.04.27.650532

**Authors:** Elita Peters, Diana A. Tanase, Lorenzo Di Michele, Roger Rubio-Sánchez

## Abstract

Biological cells use cations as signalling messengers to regulate a variety of responses. Linking cations to the functionality of synthetic membranes is thus crucial to engineering advanced biomimetic agents, such as synthetic cells. Here, we introduce bio-inspired DNA-based receptors that exploit non-canonical G-quadruplexes for cation-actuated structural and functional responses in synthetic lipid membranes. Membrane confinement grants cation-dependent control over receptor assembly and, when supplemented with hemin co-factors, their peroxidase DNAzyme activity. Cation-mediated control extends to receptor lateral distribution to localise DNA-based catalysis within phase-separated membrane domains of model synthetic cells, imitating the localisation of multimeric membrane complexes to signalling hubs in living cells. Our modular strategy paves the way for engineering from the bottom-up cation-responsive pathways for sensing, signalling, and communication in synthetic cellular systems.

## Introduction

Cell membranes possess specialised mechanisms to detect and transduce environmental cues to coordinate cellular responses. ^1^ Signal transduction pathways commonly rely on membrane receptors assembling multimeric complexes and/or undergoing conformational changes, which triggers downstream functionality through membrane-hosted reactions, e.g. (de)phosphorylation.^2^ Cations, like calcium^3,4^ and potassium^5^ ions, are central to cellular signaling, typically shuttled across biological membranes as messengers to link physicochemical stimuli to responses as varied as synaptic activation, ^6^ muscle contraction,^7^ and cellular motility.^8^

Synthetic cell science aims to build cell-like agents that replicate some of the intricate behaviours observed in living matter.^9,10^ Synthetic cells thus provide bottom-up platforms to dissect fundamental biological processes^11^ and to unlock disruptive applications in healthcare and biotechnology.^12^ By borrowing concepts and building blocks from biological and inorganic sources, solutions to engineer synthetic cells have achieved a wide range of life-like functionalities, from reconstituting cytoskeletal support^13–15^ to sustaining directional motion^16,17^ to membrane-based energy harvesting^18^ and mechanochemical sensing.^19^

Particularly promising for the construction of human-made mimics of cells are the tools of nucleic acid nanotechnology,^20,21^ as they provide robust design rules to engineer both structural and functional elements of synthetic cells. DNA and RNA nanostructures have been developed as replicas of cytoskeletal fibres^22,23^ and membrane-less organelles,^24–27^ or interfaced with lipid membranes.^28,29^ Amphiphilic DNA nanostructures have been applied to synthetic-cell membranes to program several biomimetic responses,^30–32^ ranging from molecular sensing^33^ to tissue formation^34,35^ to surface^36,37^ and transmembrane^38–40^ transport to bilayer remodelling^41,42^ and vesicular fission pathways.^43^

Given the central role of cation homeostasis in health and disease, biomimetic devices that grant structural and functional responsiveness to cations are highly desirable, as they could help realise the fundamental and applied potential of synthetic cells. However, despite recent developments, ^44–46^ pathways for embedding responsiveness to cations in bio-inspired systems, and particularly synthetic cell membranes, are still scarce. G-quadruplexes (G4s) are cation-stabilised non-canonical secondary structures, studied in biology for their role in transcriptiontranslation to regulate gene expression, telomere maintenance and disease progression^47–51^. In the context of DNA nanotechnology, G4s are convenient structural and responsive motifs^52–54^ for membrane bioengineering, as they have also been coupled to lipophilic groups to develop pathways for cation transmembrane transport^55–57^.

Here, we introduce biomimetic DNA-based membrane receptors with structural and functional responsiveness to physiologically-relevant cations. By interfacing cholesterol-modified DNA nanostructures with intermolecular G-quadruplexes, we present a membrane bioengineering strategy to modulate the formation of receptors on the surface of lipid bilayers through independent design criteria, namely the guanine content and choice of cationic con-ditions. We supplemented our cation-controlled receptors with hemin co-factors to host and regulate DNAzyme peroxidase reactions on lipid membranes. Finally, we exploit DNA receptor assembly to localise their peroxidase activity within lipid domains of phase-separated synthetic cell membranes. Our modular platform thus enables cation-actuated structural and functional responses in the membranes of synthetic cellular agents, paving the way for sophisticated biomimetic platforms towards bottomup membrane signalling, molecular transport, and division.

## Results and Discussion

### G4s assemble cation-stabilised DNA membrane receptors

Our DNA nano-devices consist of 56 base-pair (bp) long duplexes composed of four synthetic DNA oligonucleotides (Tables S1 and S2), featuring a terminal overhang with six consecutive guanines, (G)_6_, that allow for receptor assembly via an intermolecular G-quadruplex.^44^

As depicted in Fig. 1a, we produced ∼100 nm Large Unilamellar Vesicles (LUVs) with DOPC lipids and decorated their membranes with our nanostructures, which host double-cholesterol (dC) ‘anchors’ able to insert into the hydrophobic core of the bilayer.^60^ Membrane functionalisation was carried out in buffered solutions containing 1 × TE + 100 mM KCl + 87 mM sucrose, thus favouring G-quadruplex formation (see Experimental Section). Dynamic light scattering (DLS) measurements in Fig. 1b at the various LUV functionalisation stages show the expected gradual increase in hydrodynamic diameter resulting from the membrane-tethered DNA nanostructures with and without G-rich strands. When featuring G-rich strands, however, vesicle size distributions exhibit modest broadening, likely due to a low degree of LUV-LUV association mediated by intermembrane G-quadruplex formation. As seen in Fig. S1, polydispersity indices below 0.2 indicate that LUV dispersions remain predomi-nantly monodisperse with no large-scale vesicle aggregation. Note that, owing to the flexible spacer (nine thymines) separating G-runs from the double-stranded membrane anchors, G-quadruplex folding is not expected to impose significant conformational strain. This applies whether the four devices are tethered to the same membrane or on different vesicles. We therefore do not anticipate a meaningful energetic penalty or strain associated with Gquadruplex formation on the surface a single LUV. Instead, kinetic effects are likely to play a more prominent role. At the experimental concentrations of LUVs and DNA nanostructures, we expect the rate of encounter to be much higher for DNA within the same membrane than across different LUVs, owing to the much higher local concentration of membrane-bound DNA relative to LUVs in solution, despite their similar diffusion coefficients (LUVs: *D* ∼ 2–4 *µ*m^2^ s^−1^ and membrane-anchored DNA: *D* ∼ 1–5 *µ*m^2^ s^−1 61,62^). This suggests that binding rates will favour intra-LUV binding, and thus, G-quadruplexes to more likely assemble within the same vesicle membrane than across separate vesicles. We thus conclude that intra-membrane G-quadruplex formation is the predominant mode of receptor assembly.

**Figure 1.**
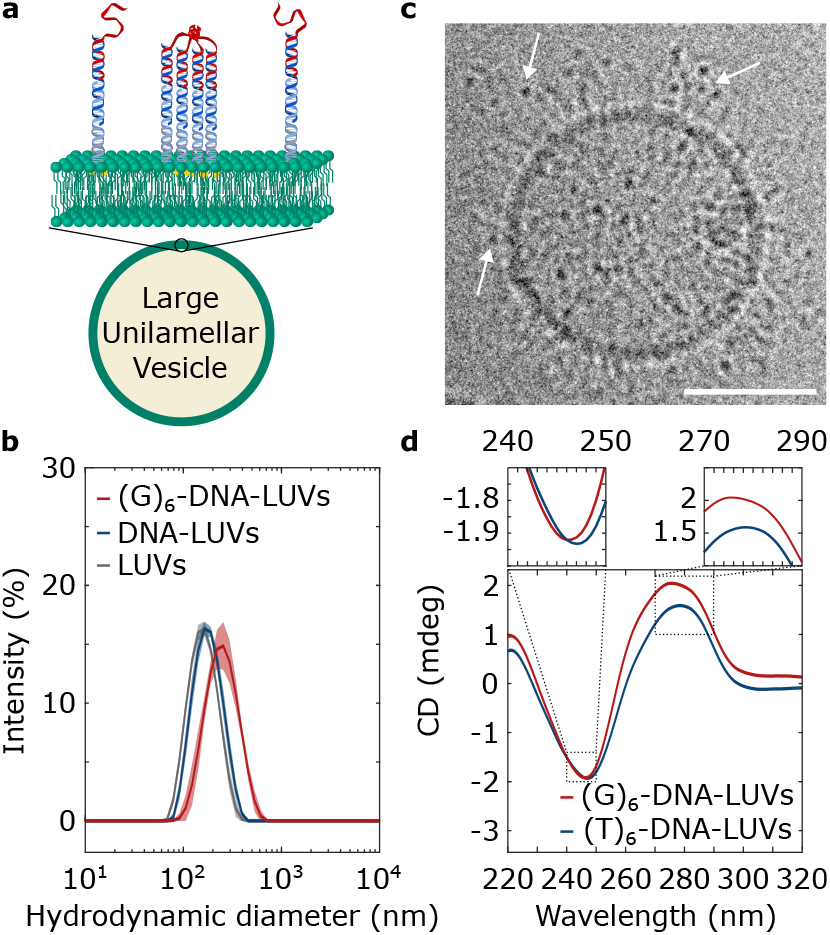
Guanine repeats in membraneanchored DNA nano-devices guide the assembly of cation-stabilised receptors through G-quadruplexes. **a** Schematic depiction of lipid bilayers functionalised with DNA nanostructures featuring a G-rich overhang that assembles into intermolecular G-quadruplexes stabilised by cations. **b** Mean hydrodynamic diameter *±* standard deviation of n = 3 measurements obtained through dynamic light scattering of Large Unilamellar Vesicles (LUVs) lacking or featuring DNA nano-devices with or without the G-rich strand that supports G-quadruplex formation. **c** Cryoelectron micrograph of representative LUV functionalised with DNA nano-devices linked to G-quadruplexes, which appear as electrondense regions^58^ (highlighted with white arrows). Scale bar = 50 nm. **d** Circular dichroism (CD) spectra of LUVs decorated with DNA nanostructures featuring either (G)_6_-overhangs (red) or a (T)_6_-repeat in place of the G-rich sequence (blue).

Cryo-electron microscopy, as seen in representative micrographs in Fig. 1c and Fig. S2, shows membrane-bound G-quadruplexes, which manifest as electron-dense regions^58^ that are absent in LUVs decorated with nanodevices lacking the G-rich strand (Fig. S3). Circular dichroism (CD) spectra of (G)_6_-DNA-decorated LUVs show peak shifts towards characteristic maximum (*λ* ≈ 263 nm) and minimum (*λ* ≈ 245 nm) wavelengths (Fig. 1d), relative to LUVs decorated with DNA nanostructures in which the G-rich sequence is replaced with a poly-thymine, (T)_6_-DNA. While the observed shift is moderate – likely due to the coexistence and different relative concentrations of G4s with double-stranded DNA anchors^63^ – the overall spectral shift and curve shape are consistent with parallel G-quadruplex topologies^64^ and with the formation of tetramolecular receptors on the membrane surface.^65^ To confirm receptor stoichiometry, we introduced a toehold-domain on the G-rich strand to enable detachment of the nano-devices from membranes with a strand displacement reaction.^66^ Following membrane functionalisation and receptor assembly, we recovered the nanostructures from LUVs and conducted a gel-shift assay, comparing their electrophoretic mobility with that of tri-, tetra- and penta-valent DNA junctions with arm lengths matching the size of a detached monomeric nanostructure. As shown in Fig. S4, receptors exhibit migration profiles similar to that of tetravalent (4-way) junctions, confirming their tetramolecular nature. The slight reduction in mobility relative to 4-way junctions is likely due to the flexible linker, which is absent from the central junction of the control multivalent nanostructures.

### Cation-controlled receptor assembly

Having assembled receptors by cross-linking individual DNA-based membrane inclusions via G-quadruplexes, we modified our nanostructure design and functionalisation conditions to control G4 formation. Specifically, we tune the length of guanine repeats and identity of the cations, both known to alter the stability of G4s.^59^

We functionalised LUVs with DNA nanodevices featuring overhang variants (G)_n=4,5,6_ (Fig. 2a) in buffered solutions with added physiologically-relevant monovalent (K^+^, Na^+^, Li^+^) or divalent (Ca^2+^, Mg^2+^) cations (see Experimental Section and Table S3 summarising cation concentration ranges in biological environments^67^). To assess G-quadruplex formation, we exploited the fluorescence en-hancement of *N* -methyl mesoporphyrin IX (NMM), which selectively binds to parallel G-tetrads. ^44,68^ Fluorescence emission spectra in Fig. S5 and S6 demonstrate that different cationic environments do not induce noticeable peak shifting. In turn, Fig. S7 shows that, relative to baseline values in our experimental buffers, NMM exhibits a moderate fluorescence increase in the presence of control (T)_6_-DNA nanostructures, likely due to weak non-specific interactions which also depend on cationic environment. As schematically depicted in Fig. 2b, we used NMM fluorimetry to compare the presence of membrane-bound Gquadruplexes with G4s assembled in the bulk from nanostructures at a nominally equal concentration ([(G)_n_] = 0.4 *µ*M, at a DNA/lipid molar ratio of ∼8 × 10^−4^). To that end and to ensure nanostructures remained fully dispersed in solution, we used non-cholesterolised DNA constructs. Cholesterol-bearing DNA nanostructures undergo micellisation in aqueous environments,^69,70^ and those would provide locally increased concentrations that would obscure the specific contributions of membrane confinement to receptor assembly. Indeed, owing to their tetrameric stoichiometry,^65^ oligonucleotide concentration is expected to have a large impact on the formation of intermolecular G4s. We therefore anticipate differences in the abundance of G-quadruplexes in the bulk relative to those tethered to LUVs, where confinement to membrane surfaces increases the effective local nanostructure concentration.

**Figure 2.**
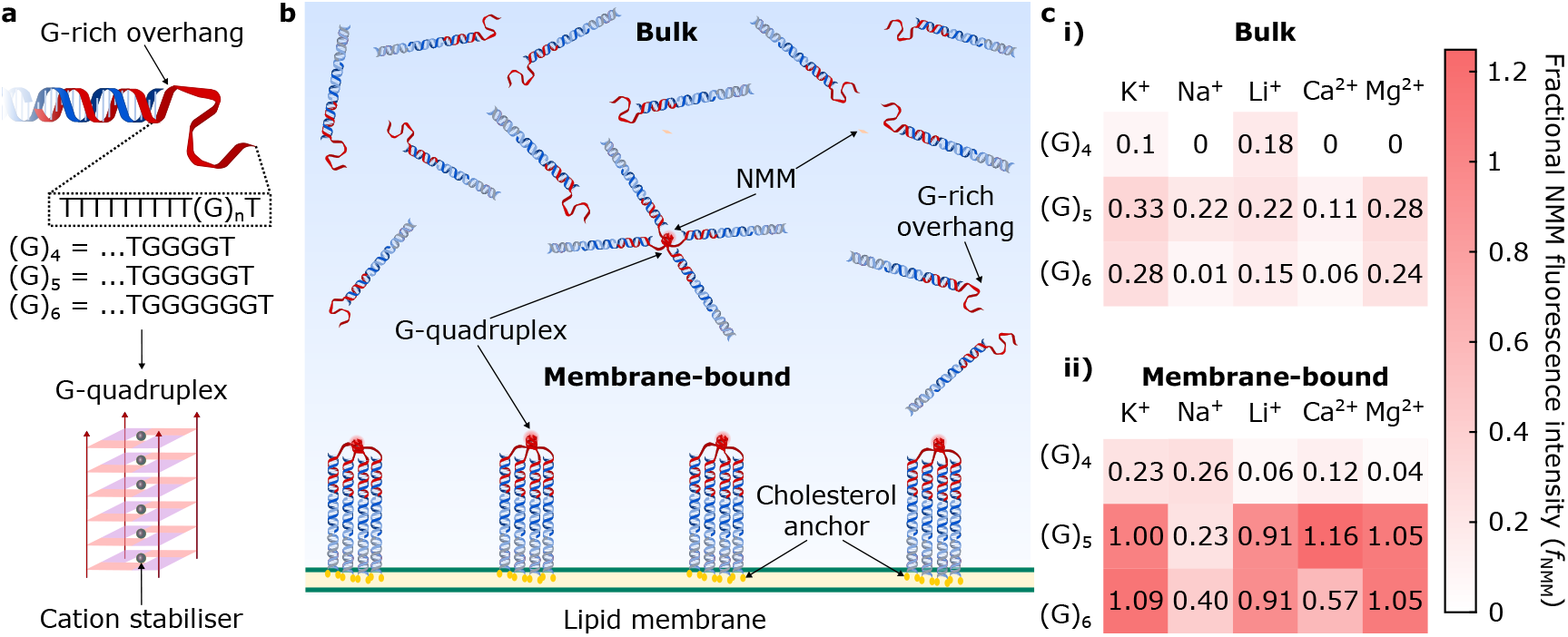
Membrane-confinement grants control over G-quadruplex formation. **a** Schematic representation of the G-rich overhang (in red) linked to DNA nano-devices, highlighting the sequence for each design variant (G_4_, G_5_, and G_6_), which can assemble into intermolecular, parallel, G-quadruplexes stabilised by cations. The schematic is illustrative, and cation coordination sites can vary (e.g. inter-planar vs in-plane) depending on cation size ^59^. **b** Graphical depiction of tetramolecular receptors assembled through G-quadruplexes in the bulk and when membranebound via double-cholesterol anchors. Parallel G-tetrads selectively bind NMM and enhance its fluorescence, enabling to monitor the abundance of G-quadruplexes. **c** Fractional NMM fluorescence intensity (*f*_NMM_) heatmaps (see Fig. S11 for the associated standard deviation *δf*_NMM_) summarising the relative abundances of receptors in: i) the bulk and ii) membrane-bound as a function of monovalent (K^+^, Na^+^, or Li^+^) and divalent (Ca^2+^ or Mg^2+^) cations for designs G_4_, G_5_, and G_6_ (at a nominal concentration of [(G_n_)] = 0.4 *µ*M).

To quantify the relative abundance of G-quadruplexes, we compute a fractional NMM fluorescence intensity (*f*_NMM_), defined as the cation-specific background-subtracted average intensity of DNA-decorated LUVs normalised by the NMM fluorescence intensity of (G)_5_-DNA-LUVs in K^+^ (see Experimental Section). The latter was chosen as it is the condition in which we found the highest fraction of G4s when nanostructures were allowed to assemble in the bulk via thermal annealing at high concentrations ([(G)_n_] = 6 *µ*M), as quantified with agarose gel electrophoresis (Fig. S8) and observed with NMM fluorimetry (Fig. S9). Note that representative, time-invariant, fluo-rescent profiles in Fig. S10 confirm that both G-quadruplex formation and NMM stacking onto G-tetrads have reached thermodynamic equilibrium.

When the nanostructures were dispersed in the bulk, *f*_NMM_ values ≤0.33 shown in Fig. 2c-i (and their comparatively high standard deviation, *δf*_NMM_, in Fig. S11) suggest low G4 abundance relative to DNA-decorated LUVs. Indeed, tethering our nano-devices to lipid membranes, and thus increasing their effective local concentration, led to systematically higher *f*_NMM_ values across the tested G-run lengths and cationic conditions (Fig. 2c-ii), confirming the expected effect on G4 assembly. The shortest G-repeat, (G)_4_, with low *f*_NMM_ values – albeit higher than their bulk counterparts (with the exception of Li^+^) – has relatively low G-quadruplex abundances compared to LUVs decorated with (G)_5_ in KCl, where *f*_NMM_ ≡ 1. CD signatures of (G)_4_-decorated LUVs in K^+^ show slight characteristic peak shifts (Fig. S12), consistent with the parallel quadruplex topologies observed for LUVs carrying (G)_6_-(Fig.1c) and (G)_5_-overhangs (Fig. S13). Note that, while the combination of CD spectroscopy, NMM fluorimetry, and gel-shift assays indicate the formation of parallel, tetramolecular, G-quadruplexes, the number of stacked tetrads in G4s cannot be unambigously assigned without high-resolution crystallographic data. We speculate, however, that tier number should be equal to the length of the G-run^71^.

Various cationic compositions, namely Li^+^ and Mg^2+^, enable G4 formation in (G)_5_ and (G)_6_ constructs to a comparable degree to that of (G)_5_-membranes with K^+^ (reaching *f*_NMM_ ∼ 1 and within the experimental error in Fig. S11). Similarly, (G)_5_-DNA-decorated LUVs in Ca^2+^ have *f*_NMM_ values moderately higher than 1, suggesting favourable G4 formation. Interestingly, (G)_6_- and (G)_5_-runs in the presence of Na^+^ show the lowest relative abundance, with *f*_NMM_ ∼0.40 and *f*_NMM_ ∼0.23, respectively.

To substantiate our hypothesis that the higher local DNA concentrations achieved with membrane confinement influence G-quadruplex formation and stability, we performed simple numerical calculations using a thermodynamic description of tetramolecular G4 assembly from four monomeric DNA constructs (see Supplementary Note I). Briefly, we relate the fraction of DNA in G4 constructs (*p*) with the total DNA concentration (*C*) and the standard free energy of assembly (Δ*G*^*°*^), capturing the sharp changes of *p* with increasing DNA concentrations (Fig. S14). As an example, in the case of Δ*G*^*°*^ = −30*k*_B_*T*, low DNA concentrations, such as those used in our bulk assembly assays, yield negligible probabilities of G4 assembly *p*. In contrast, when membrane-confined, the estimated local concentration (∼171 *µ*M) yields *p* ∼ 0.7, corresponding to a 14-fold increase in G-quadruplex formation probability.

While the specific free energies of G4 formation in our experiments are not known, the numerical solutions indicate that the high local DNA concentrations on membranes can substantially increase the probability of G-quadruplex assembly, even with ions that, like lithium, do not stabilise G4s in the bulk. Thus, our calculations provide a thermodynamic rationale for how membrane attachment, by increasing the local concentration of DNA, shifts the equilibrium in favour of G4 assembly even in the presence of weakly coordinating cations. Notably, molecular dynamic simulations show that Li^+^ can indeed weakly coordinate with G-tetrads, albeit with free energies of bond formation ∼15 kcal/mol less favourable than K^+^ ions.^72^ We thus argue that, despite weak coordination of Li^+^, under membrane confinement – where DNA is locally concentrated – G-quadruplex formation probabilities would be substantially higher.

Furthermore, in view of the higher G4-stabilisation expected to occur with Na^+^ cations compared to Mg^2+^ or Li^+^, the trends on G4 stability seen for DNA-LUVs (Fig. 2c-ii) could also be influenced by the interplay between DNA-DNA and DNA-lipid interactions. Indeed, each cation species has a distinct affinity for phosphate groups present on both DNA and PC lipid headgroups. Cations have been shown on DNA-lipid systems to screen

Coulomb interactions and/or mediate bridging to different extents, thus inducing differences on the membrane attachment of DNA nanostructures.^45^ It is thus possible, for instance, that divalent cations further increase nanostructure effective concentration (and favour G4 formation) by bridging membranes with DNA duplexes.^45,46^

To gain insight on the assembly dynamics of our receptors, we decorated LUV membranes with nano-devices in the presence of KCl but lacking the G-rich strand, thus preventing G4s. We subsequently added a mixture of stoichiometrically-adjusted (G)_6_- or (G)_5_-strands and NMM, allowing oligonucleotides to rapidly diffuse, hybridise with membrane-bound nanostructures, and assemble into K^+^-stabilised G-quadruplexes where NMM could stack, leading to fluorescence enhancement (see Experimental Section). Consistent with the high local concentrations of DNA nanostructures upon membrane-confinement (∼171 *µ*M), NMM fluorescent profiles in Figs. S15 and S16 suggest fast G-quadruplex assembly. Equilibration of fluorescence under 250 seconds for both design variants is likely limited by the diffusion of single-stranded oligonucleotides throughout the samples, as we expect the diffusion timescales of free oligonucleotides in solution (at the relevant concentrations and lengths)^73^ to be on the order of hundreds of seconds.

### Cation-actuated DNA receptors with membrane-hosted catalytic activity

Our platform to control the formation of cation-stabilised receptors can be linked to the functionality of DNAzymes underpinned by G4s. A prominent example is that of the horseradish peroxidase (HRP)-mimicking DNAzyme, composed of hemin co-factors stacked onto Gtetrads.^74^ Binding to G4s strongly enhances the catalytic activity of hemin when converting AmplexRed (AR) to fluorescent resorufin in the presence of hydrogen peroxide (H_2_O_2_).^75^

We thus produced LUVs decorated with our nano-devices and incubated them with hemin co-factors, resulting in receptor assembly hosting the HRP-mimicking DNAzyme (Fig. 3a). As shown in Fig. 3b and Figs. S17 and S18, we monitored resorufin production by means of fluorescence spectroscopy. In view of the trends summarised in Fig. 2, we expect our various designs and cationic conditions to result in different amounts of receptors available for hemin to bind, and thus the various conditions to produce differences in peroxidation rates.

**Figure 3.**
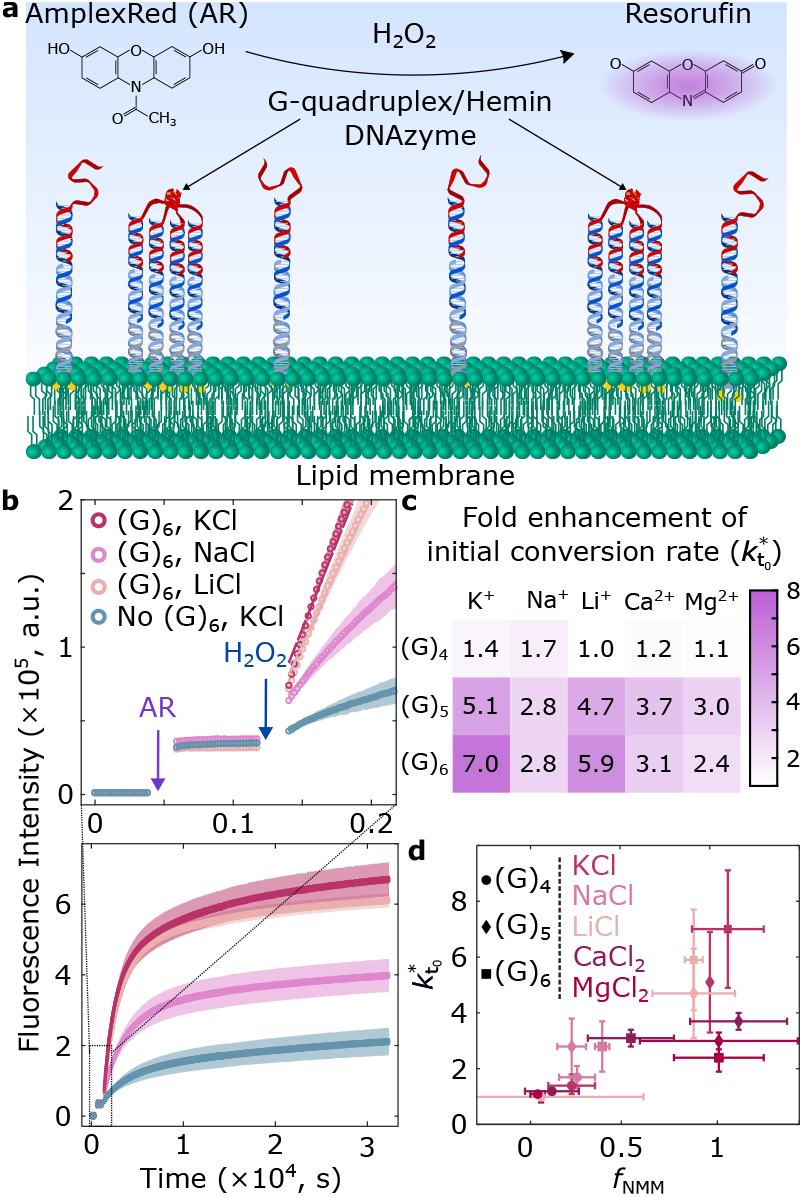
Cation-controlled DNA receptors for membrane-hosted peroxidase activity. **a** Schematic showing the incorporation of hemin co-factors to form the horseradish peroxidase mimicking DNAzyme, which in the presence of hydrogen peroxide (H_2_O_2_) catalyses the conversion of AmplexRed (AR) to fluorescent resorufin. **b** Representative resorufin fluorescence intensity profiles of DNA-decorated LUVs with and without (G)_6_-rich strands and added hemin co-factor in KCl, NaCl, and LiCl. Inset shows, at early times, the fluorescence intensity of LUVs before/after AR addition, and its increase upon initiating the reaction with H_2_O_2_. Solid lines are linear fits used to extract initial reaction rates 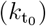. **c** Heatmap of fold change in initial reaction rate 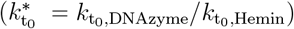 for receptors asssembled from either (G)_n=4,5,6_ as a function of cationic conditions. **d** Rate fold changes 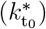 correlate with receptor relative abundance (*f*_NMM_); statistical significance assessed via Spearman test: *ρ* = 0.7876, *p*−value = 4.89 *×* 10^−4^.

To quantify the influence of our assembly platform on resorufin production, we extracted the initial reaction rates 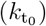 from linear fitting of the fluorescent traces (see inset Fig. 3b and Methods) and computed their fold enhancement relative to DNA-LUVs (i.e. lacking G4s) in the presence of hemin co-factors using 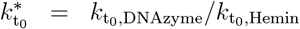. To rule out the possibility that non-specific interactions between hemin with duplexes or unstructured DNA could contribute to catalysis, we compared peroxidation across (G)_6_-DNA-LUVs, (T)_6_-DNA-LUVs, DNA-LUVs, and non-functionalised LUVs. As shown in Fig. S19, control conditions exhibit largely overlapping behaviours, confirming that, in our platform, hemin catalytic activity is enhanced in the presence of G-quadruplexes.

The heatmap in Fig. 3c (and the associated propagated uncertainty 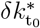 in Fig. S20) readily confirms that various conversion rates can be accessed by design. Note that, consistent with our observations on relative G4 abundance in Fig. 2c)ii, LUVs functionalised in the presence of LiCl result in higher 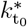 values compared to NaCl. We thus further explored the relationship between relative receptor abundance and fold-change in initial peroxidation rate 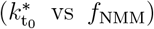. As shown in Fig. 3d, we find a statistically-significant correlation (non-parametric statistical correlation Spearman test: *ρ* = 0.7876; *p™*value = 4.89 × 10^−4^), with greater fold-changes corresponding to higher *f*_NMM_ values. Small deviations to this trend emerge for specific conditions with divalent cations which, despite reaching high *f*_NMM_ values, exhibit moderately lower fold-changes in peroxidation rate. We ascribe this decrease to a lower catalytic efficiency of hemin in the presence of divalent cations, as seen in the fluorescent traces on Figs. S17 and S18, where peroxidation is catalysed solely by the co-factor. Importantly, although minor inter-LUV asso-ciation was observed with DLS in Fig. 1b and Fig. S1, we do not expect clustering to substantially alter DNAzyme functional performance given that hemin co-factors and the reaction substrates (AmplexRed and H_2_O_2_) are small molecules that could still freely diffuse through LUV-LUV contacts.

### Localising receptor activity in membrane domains of synthetic cells

The cation-dependent assembly and activity of our DNA receptors can be readily coupled to membrane phase separation. Their synergy enables to mimic the re-organisation and activity of multimeric protein nanomachines within cellular membrane domains.

To exemplify the applicability of our platform for synthetic-cell engineering, we produced Giant Unilamellar Vesicles (GUVs) as the chassis of our model synthetic cells, and decorated their membrane with our receptors. Synthetic cells, prepared from lipid mixtures containing DOPC/DPPC/Cholestanol in 2:2:1 molar ratios, displayed membrane phase separation at room temperature with co-existing liquid-ordered (*L*_o_) and liquid-disordered (*L*_d_) domains.^77^ A fluorescent TexasRed-DHPE lipid marker was included to stain *L*_d_ phases, while our DNA nano-devices were modified to include a fluorescein probe to monitor their lateral organisation by means of confocal microscopy.

Given the partitioning of double-cholesterol anchors to lipid domains in phase-separated membranes,^36,78,79^ DNA receptor assembly enriched *L*_o_ domains by cross-linking four sets of dC anchors, as shown schematically in Fig. 4a and with representative confocal micrographs in Fig. 4b. Segmentation of confocal equatorial micrographs allowed us to assess DNA nanostructure partitioning by comparing average fluorescence intensities in *L*_o_ and *L*_d_ phases (see Supplementary Note III). Expectedly, as seen in Fig. S21, monomeric DNA nanodevices (i.e. lacking the G-rich strand) moderately accumulate in *L*_o_ domains. Subtle differences in their *L*_o_-partitioning tendencies already emerge between added salts, further supporting our hypothesis of cations modulating electrostatic DNA-DNA and DNA-lipid interactions. Stronger partitioning of dC-anchored nanostructures is achieved in the presence of di-valent cations, while Li^+^ promotes greater *L*_o_-accumulation when compared to Na^+^ and K^+^. These differences can be ascribed to variations in screening effectively the negatively-charged phosphate groups present both on the DNA backbone and the lipid headgroups on the membrane surface^45^.

**Figure 4.**
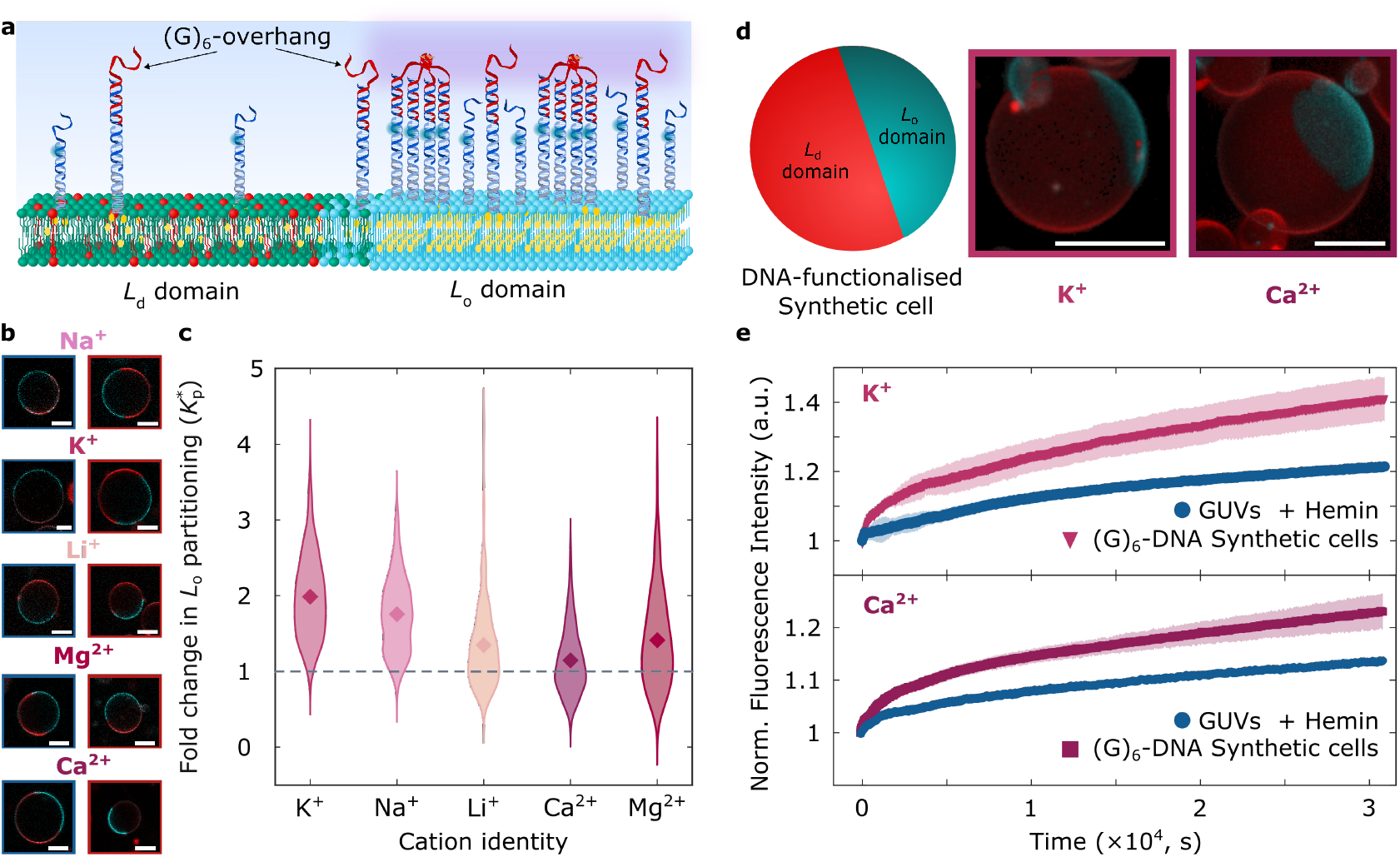
Cation-dependent assembly localises receptors and their peroxidase activity in lipid domains of synthetic cell membranes. **a** Schematic of the organisation in lipid domains of DNA nano-devices with and without (G)_6_-strands, and therefore, the possibility of receptor assembly localising peroxidase activity when supplemented with hemin co-factors. **b** Representative confocal equatorial micrographs of GUVs functionalised with DNA nano-devices lacking (left, framed in blue) and featuring (right, framed in red) (G)_6_-strands to support receptor formation in buffered solutions with added monovalent (Na^+^, K^+^, or Li^+^) or divalent (Mg^2+^, or Ca^2+^) salts. The TexasRed-DHPE signal, staining the *L*_d_ phase, is shown in red, while that of fluorescein (FAM)-labelled DNA nanostructures is shown in cyan. **c** Violin plots of the fold change in *L*_o_-partitioning 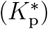, computed from confocal micrographs (see Experimental Section) of DNA-decorated GUVs in either monovalent (Na^+^, K^+^, or Li^+^) or divalent (Mg^2+^, or Ca^2+^) cationic conditions, showing the cation-dependent membrane distribution of DNA receptors. Diamonds are mean fold change values, while the dashed line denotes no change 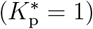. **d** Schematic depiction of a DNA-decorated synthetic cell (left) and 3D views of reconstructions (obtained from Volume Viewer, FIJI^76^) from confocal z-stacks of representative DNA-decorated GUVs (in the presence of K^+^ or Ca^2+^) as synthetic cell models with *L*_o_-localised functionality. **e** Normalised fluorescence profiles monitoring resorufin production from DNA-functionalised synthetic cells in the presence of K^+^ (top) or Ca^2+^ (bottom) relative to GUVs lacking DNA functionalisation but supplemented with the hemin co-factor, showing the possibility to localise peroxidase activity in lipid domains of synthetic cell membranes. All scale bars = 10 *µ*m.

Conversely, when nanostructures feature the (G)_6_-overhang, thus enabling G4 formation, devices systematically show stronger partitioning relative to their state prior to the addition of the G-rich strand. In Fig. 4c we summarise the fold enhacement in *L*_o_-partitioning 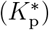 of (G)_6_-DNA membranes against that of functionalized GUVs lacking (G)_6_-overhangs. Fold changes, with statistically significant values of 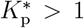(*p <* 1.2 × 10^−16^; one-tailed non-parametric Wilcoxon signed-ranked test – see Table S4 for individual *p*−values) are indicative of stronger affinities for *L*_o_ domains due to the increased number of cholesterol anchors per nanostruture.^36^ The latter therefore supports the presence of tetramolecular receptors and highlights our ability to regulate their membrane distribution in a cation-dependent fashion. Notably, due to the macroscopic sizes of GUVs, membrane curvature effects in domain partitioning are expected to be negligible, as the membrane can be considered to be locally flat relative to the size of the nano-devices. This interpretation is further supported by the absence of noticeable correlation between domain accumulation and vesicle size across different cationic compositions (Fig. S22 and Table S5 with non-parametric Spearman *ρ* and *p*−values).

We subsequently sought to localise receptor peroxidase activity in *L*_o_ domains of our synthetic cells. As proof-of-concept, we selected K^+^ or Ca^2+^ as monovalent or divalent cation messengers, respectively, and incubated DNA-decorated synthetic cells (shown in Fig. 4d and Fig. S23 with representative 3D views from confocal z-stacks) with hemin co-factors. We monitored resorufin production upon triggering synthetic cell activity with H_2_O_2_, as summarised in Fig. 4e with normalised fluorescent profiles (as well as on Figs. S24 and S25 with individual replicate profiles), where expectedly synthetic-cell membranes featuring our receptors exhibit faster and higher conversion than membranes lacking functionalisation in the presence of the hemin co-factor. Epi-fluorescence micrographs in Fig. S26 of synthetic cells after peroxidation confirm their stability throughout the experimental timescales. While direct imaging of resorufin being produced at lipid domains is limited by diffusion timescales, our results showing the preferential accumulation of membrane-bound, catalytically-active, receptors in liquid-ordered phases strongly suggest that catalytic activity is predominantly localised in *L*_o_-domains. Therefore, with our platform we show the possibility to localise and regulate activity within membrane domains of synthetic cells using different cation messengers.

## Conclusions and Outlook

In summary, we have introduced biomimetic receptors with cation-controlled assembly, catalytic activity, and distribution in synthetic cell membranes. Our modular platform affords stimulus-responsive assembly of catalytically-active receptors that sustain lateral re-shuffling on membrane surfaces, integrating cation-responsiveness with spatial organisation and catalytic function. Our tetrameric membrane receptors exploit the formation of intermolecular G-quadruplexes to assemble on the surface of lipid bilayers. By exploiting the increased effective local concentrations induced by membrane confinement, we show that receptor formation can be controlled with different cations. Similarly, cation-mediated control allows us to regulate the rate of membrane-hosted reactions as exemplified with peroxidation using the HRP-mimicking DNAzyme. Finally, by interfacing our cation-actuated receptors with phase-separated membranes in GUVs, we established a link between the presence of cations and the lateral organisation of functional DNA-based membrane inclusions. We applied our nano-devices to host peroxidase activity within lipid domains of model synthetic cells, thereby imitating the ability of cells to localise reactions within cell-membrane domains as hypothesised to occur during signalling cascades. ^80,81^

Our strategy has direct applications in bottom-up synthetic biology to engineer advanced synthetic cell models. Because of its modularity and the versatility of DNA-nanotechnology, our cation-responsive approach to crosslink individual membrane inclusions can be readily adapted to guide the assembly of more intricate DNA and RNA origami nano-devices. Indeed, simple design changes can enable coupling the presence of cations to the formation of cortex-like platforms^82,83^ and membrane-remodelling nanostructures,^42,43^ unlocking the development of cation-dependent pathways for cellular motion, trafficking, division, and bio-inspired synaptic transmission. Similarly, the ionic environments assessed in our work are within cation concentration ranges relevant to in vitro transcription and cell-free expression systems, ^84,85^ suggesting possible synergies with the functionality of our receptors for synthetic cell engineering. In addition, the possibility of destabilising G-quadruplexes (e.g. with photo-induced damage^86,87^ or chelating agents^88^) paves the way for the development of sophisticated dynamic behaviours that respond to both physical and chemical stimuli.

Similarly, our DNA receptors could underpin the investigation of fundamental biological processes. One can envision our nano-devices as probes^89^ tethered to cellular membranes that could report on changes in electrochemical activity, allowing, for instance, monitoring the kinetics of cation waves^90,91^ as well as cation transport across biomembranes.^92–94^

Finally, the effect of surface confinement on local concentration, as exploited here to regulate peroxidase activity, could be leveraged to influence the action of other DNAzymes, ribozymes^95^ and (split) aptamers,^96^ many of which are typically underpinned by G4s. For instance, it could be possible to finely tune the efficiency of Mg-dependent cleaving nanostructures^97^ or the allosteric modulation of enzy-matic action,^98^ thereby unlocking the possibility to host in (bio)membranes a wider range of reactions useful in biosensing, biotechnology, and bioengineering.

## Acknowledgement

E.P. acknowledges funding from the NanoDTC (EP/S022953/1) through the NanoFutures Scholars programme. R.R.S. acknowledges funding from the Biotechnology and Biological Sciences Research Council through a BBSRC Discovery Fellowship (BB/X010228/1) and from Wolfson College, Cambridge. L.D.M. and D.A.T. acknowledge support from the European Research Council (ERC) under the Horizon 2020 research and innovation programme (ERC-STG No 851667 - NANOCELL). L.D.M. also acknowledges support from a Royal Society University Research Fellowship (UF160152, URF \R\ 221009). The authors acknowledge assistance from the CryoEM facility (D. Chirgadaze and L. Cooper) in the Department of Biochemistry, University of Cambridge. A dataset in support of this work can be accessed free of charge at: https://doi.org/10.17863/CAM.120318

## Supporting Information Available

The Supporting Information is available at:

The following file is available free of charge including:

- Experimental methods, supplementary notes, supplementary figures, supplementary table with cation concentrations in physiological environments, supplementary tables with statistical significance and correlation values, and DNA sequences of the nanostructures used throughout this work.

## TOC Graphic

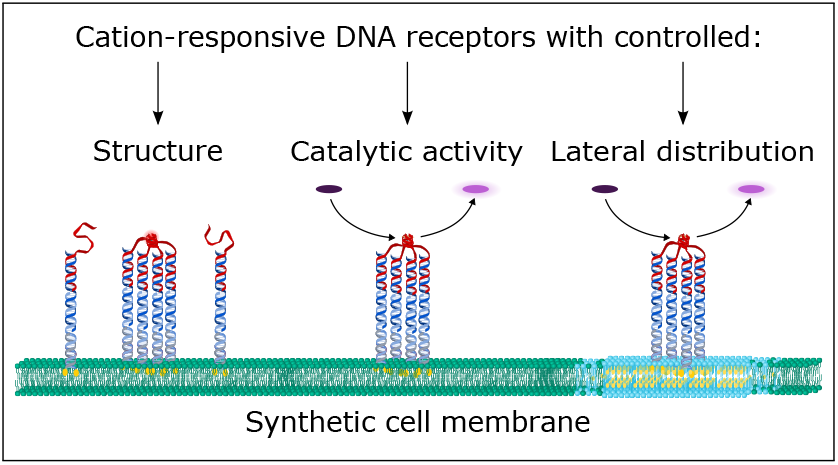

