## Supporting Information for "Cation-controlled assembly, activity, and organisation of biomimetic DNA receptors in synthetic cell membranes"

#### Experimental Methods

##### Extrusion of Large Unilamellar Vesicles

Large Unilamellar Vesicles (LUVs) were prepared via extrusion as described in previous contributions.[1,2] Briefly, glass vials were cleaned following a sonication routine with cycles of 15 minutes (2% Hellmanax III [HellmaAnalytics], isopropanol, and milli-Q water). After drying, 150 $\mu$L of 1,2-dioleoyl-sn-glycero-3-phosphocholine (DOPC, Avanti Polar Lipids)

lipids (dispersed in chloroform at 25 mg $mL^{-1}$) were pipetted into the clean 1.5 mL glass vials and placed in a dry silica desiccator under vacuum for at least an hour; the lipids were rehydrated in solutions containing 300 mM sucrose by gentle vortexing for approximately 5 minutes. Resuspended vesicles then followed a 5× freeze-thaw cycle with liquid nitrogen to promote unilamellarity. Using an extrusion kit (Avanti Polar Lipids) assembled with Whatman filters and membranes (with pore size of 0.1 $\mu$m) purchased from Fisher Scientific, the resuspended vesicles were pushed through the set-up 31 times to improve monodispersity. Vesicles were stored at room temperature.

#### Electroformation of Phase-separating Giant Unilamellar Vesicles

Giant Unilamellar Vesicles, prepared in a 300 mM sucrose solutions, were produced via electroformation as done previously.[3–6] Lipid mixtures contained DOPC (chain melting temperature -17°C), 1,2-dipalmitoyl-sn-glycero-3-phosphocholine (DPPC, chain melting temperature 41°C; Avanti Polar Lipids), and cholestanol (Sigma-Aldrich) at 2:2:1 molar ratios. Mixtures were doped with fluorescent TexasRed-DHPE (Invitrogen) at 0.8% molar ratio to preferentially stain $L_{\mathrm{d}}$ membrane domains. Subsequently, 45 $\mu$L of lipid mixtures (4 mg $mL^{-1}$) were gently spread on the conductive side of clean indium tin oxide (ITO) slides at 60°C. Slides were then placed under vacuum in a dry silica dessicator for ∼1 hour and subsequently assembled into an electroformation chamber using a ∼1 mm thick polydimethylsiloxane (PDMS) spacer enclosing approximately 200 $\mu$L of degassed sucrose buffer. Electroformation chambers, placed inside a pre-heated oven at $T > 60$°C and connected to a frequency generator via clamps, were subjected to a sinusoidal alternating current (AC) with voltage amplitude of 2 V. Electroformation was carried out at a frequency of 10 Hz for 2 hrs and then followed by 2 Hz for 1 hr. GUVs were retrieved and stored at room temperature in the dark to minimise photo-bleaching and photo-oxidation.

#### Design and assembly of DNA nanostructures

DNA nano-devices, designed using the NUPACK suite,[7] are based on previously reported nano-devices.[5,8] DNA oligonucleotides were purchased lyophilised (Integrated DNA Technologies [IDT] and Eurogentec), and were reconstituted to a nominal concentration of 100 $\mu$M in Tris – Ethylenediaminetetraacetic acid (EDTA) buffer (1× TE: 10 mM Tris + 1 mM EDTA, pH 8.0). The strands composing the DNA nano-devices were mixed at stoichimetric ratios and subjected to a slow quenching temperature ramp on an Alpha Cycler (PCR Max) thermal cycler (95°C down to 4°C at a rate of –0.5°C min $^{-1}$) in buffered saline solutions, containing either 1× TE + 100 mM monovalent salts (NaCl, KCl, or LiCl) or 1× TE + 2 mM divalent salts ($MgCl_2$ or $CaCl_2$).

#### Membrane Functionalisation with DNA nanostructures

Pre-assembled DNA nano-devices were incubated overnight with vesicles, either LUVs or GUVs (16.7 $\mu$L of DNA solution + 9.2 $\mu$L vesicle solution, in 57.4 $\mu$L of buffer resulting in an iso-osmolar mixture).[5,6,9] ”Following overnight incubation, 8.35 $\mu$L of stoichiometrically-adjusted G-rich strands (dispersed in 1× TE) were mixed with a second correction buffer (10 $\mu$L) to produce osmolarity and ionic strength equal to samples containing pre-functionalised GUVs. The mixture was then gently added to the vesicles already decorated with DNA nano-devices, resulting in an osmotically-balanced solution containing either 1×TE + 87 mM of sugar (sucrose for LUVs and glucose for GUVs) + 100 mM of monovalent salt or 1×TE + 281 mM of sugar + 2 mM divalent salt, as required. Samples were then left under rotation overnight to allow for the G-rich strands to diffuse and hybridise the membrane-anchored dC modules.

The concentrations we chose to work with for NaCl, KCl, $MgCl_2$, and $CaCl_2$ are within relevant concentration ranges found in biological environments. In Table S3 we summarise the typical concentrations for each cation species in both intra- and extra-cellular media.[10]

We note, however, that the concentrations of $Li^+$ we used are not physiologically relevant. We have decided to work at this concentration to compare G-quadruplex assembly with that occurring in the presence of other monovalent cations at equal concentrations.

#### Dynamic Light Scattering (DLS)

DLS measurements of DNA-decorated LUVs were performed using a Zetasizer Nano ZSP (Malvern Panalytical) with an excitation wavelength of $\lambda = 633\,\text{nm}$ and a scattering angle fixed at 173°. To that end, $100\,\mu$L of each sample were loaded into a low-volume quartz cuvette. For each condition, n = 3 measurements, which themselves consisted of 12 runs, were collected.

#### Circular Dichroism (CD)

Circular Dichroism spectra were acquired a J-1500 CD Spectrometer (JASCO Corporation). Briefly, $150\,\mu$L of each sample were pipetted into a 1 mm path-length quartz cuvette. Spectra were acquired between 190 and 350 nm, with a 0.5 nm bandwidth and a 1 nm data pitch. Ten spectra were accumulated for each condition and automatically averaged by the spectropolarimeter software. Data processing included blank correction by subtracting the spectrum of LUVs dispersed in saline buffer with added sucrose (1×TE + 100 mM KCl + 87 mM sucrose). All plotted spectra were processed in the spectropolarimeter software via smoothing through a Savitzky-Golay routine with a convolution width of 17.

#### Agarose Gel Electrophoresis (AGE)

Agarose gels were prepared at 1.5% (w/v) in Tris-Borate-EDTA (TBE, Sigma-Aldrich, 89 mM Tris-borate, 2 mM EDTA, pH 8.3) buffer. The mixture was dissolved with pulsed-heating using a microwave. Subsequently, SYBR Safe DNA gel stain (Invitrogen) was added at 0.1% (v/v), and mixed through gentle swirling. Agarose was then casted to a thickness of approximately 5 mm. and allowed to set for 1 hour. The gel was subsquently placed in an electrophoresis chamber and covered with TBE. 15 $\mu$L of annealed samples were loaded onto the gel alongside a DNA reference ladder (1Kb bp GeneRuler, Thermo Scientific). A potential of 70 V (3.75 V $cm^{-1}$) was applied for 60 minutes. Finally, the gel was visualised with a G:BOX Chemi XX6 (SYNGENE) imager.

In conditions where G-quadruplex assembly is favoured, we observed the presence of two bands: i) one with higher electrophoretic mobility and thus migrating further in the gel, corresponding to monomeric nano-devices, and ii) a band with lower mobility, which corresponds to tetramolecular nanostructures stabilised by G-quadruplexes. To quantitatively assess the assembly tendencies, micrographs were loaded onto a custom-built pipeline to sample and extract the intensity profiles for each lane. From each profile containing two bands, the area under the intensity profile for each band was computed numerically to estimate a fractional intensity using:

$$f = \frac{A_{\text{low}}}{A_{\text{low}} + A_{\text{high}}} \tag{S1}$$

where $A_{\text{low}}$ and $A_{\text{high}}$ are, respectively, the areas corresponding to the bands with low and high electrophoretic mobility. Fractional values $f$, which are proportional to the probabilities of finding a nanostructure in either the monomeric or tetramolecular state, were then used to compute the heatmap in Fig. S3.

#### Cryogenic Electron Microscopy (Cryo-EM)

Cryo-EM was used to visualise G-quadruplexes on the surface of lipid membranes. Briefly, Quantifoil 1.2/1.3 300 Cu mesh grids were double glow discharged, once per side with 25mA for 60 seconds. Subsequently, 3 $\mu$L of DNA-decorated LUVs were loaded onto grids in a Vitrobot 1 (Thermo Fisher Scientific) set to 4°C, 95% humidity, using 3-second blot times and -5 blot force, and then plunged into liquid ethane. Grids were clipped and transferred using liquid nitrogen. Micrographs were acquired using a Talos Arctica transmission electron microscope (Thermo Fisher Scientific) operated at 200 kV equipped with a Falcon 3 detector. Grid preparation and micrograph acquisition were performed at the CryoEM facility in the Department of Biochemistry, University of Cambridge.

#### Fluorimetry

Fluorescence spectroscopy was carried out using a BMG CLARIOstar Plus microplate reader using Clear-bottom, non-treated, 384 well-plates (NUNC).

##### Monitoring degree of receptor assembly with NMM fluorimetry

To assess the extent of G-quadruplex formation, 8.35 $\mu$L of stoichiometrically-adjusted NMM were mixed with 10 $\mu$L of our second correction buffer resulting in osmotically-balanced solutions containing either 1×TE + 87 mM of sugar (sucrose for LUVs and glucose for GUVs) + 100 mM of monovalent salt or 1×TE + 281 mM of sugar + 2 mM divalent salt, as required. The mixture was gently added to DNA-decorated LUVs, prepared as detailed in "Membrane Functionalisation with DNA nanostructures" and incubated for at least 30 minutes. Samples were then pipetted into 384 well-plates, covered with MicroAmp Optical Adhesive Film (ThermoFisher Scientific) to prevent evaporation, and allowed to equilibrate and thermalise ($T$ = 25°C) inside the microplate reader for at least an hour. Measurements were performed using $\lambda_{\text{excitation}} = 388 \pm 15$ nm and $\lambda_{\text{emission}} = 670 \pm 15$ nm with fixed gains and focal heights, adjusted by the instrument software in each independent repeat. Individual measurements, such as those shown in Fig. S5 with representative fluorescent profiles, consisted of acquisition for 30 cycles – each composed of 30 flashes per well with orbital averaging scanning mode (2 mm of diameter) across 30 flashes. The collected traces were then averaged to compute $f_{\mathrm{NMM}}$ using

$$f_{\mathrm{NMM}} = \frac{I_{\mathrm{NMM,x}} - I_{\mathrm{background}}}{I_{\mathrm{NMM,Max}} - I_{\mathrm{background}}} \tag{S2}$$

where $I_{\mathrm{NMM,x}}$ is the fluorescence intensity of NMM in a given design variant and cationic condition and $I_{\mathrm{background}}$ is the signal of NMM when dispersed in the respective buffered saline solution supplemented with sucrose in the presence of non-cholesterolised $(T)_6$-DNA nanostructures, allowing to account for NMM non-specific fluorescence changes in each cationic environment (Fig. S7). In turn, $I_{\mathrm{NMM,Max}}$ is the intensity of $(G)_5$-DNA-LUVs in KCl. We chose to normalise relative to this condition since $(G)_5$ in KCl exhibited the highest amount of G4s when non-cholesterolised nanostructures were assembled and allowed to equilibrate via thermal annealing, as seen in Figs. S8 and S9. Values and their errors (propagated from n = 6 repeats) were used to construct the heatmaps in Figs. 2 (main text), S9, and S11 to summarise the relative G4 abundances across the explored parameter space.

NMM emission spectra (Figs. S5 and S6) were acquired exciting at $\lambda_{\mathrm{excitation}} = 388 \pm 15$ nm, with collection from $\lambda = 550$ to $720$ nm.

##### Assessing the kinetics of receptor assembly with NMM fluorimetry

To assess the formation kinetics of our cation-stabilised nanostructures, the experimental approach consisted of adding both NMM and the G-rich strand (dispersed in 1×TE without added salts to prevent assembly) to LUVs decorated with DNA nanostructures. In this manner, G-rich strands could diffuse and hybridise the nano-devices on the membrane surface, where G-quadruplexes could then assemble and mediate the fluorescence enhancement of NMM molecules. This approach was necessary given that cations are required for membrane-attachement of DNA nanostructures,[1,2] and therefore our DNA nano-devices would not bind to lipid membranes in conditions devoid of cations. Thus, after overnight incubation, DNA-decorated LUVs were mixed with a correction buffer such that the addition of 5 $\mu$L containing NMM + G-rich strands would result in iso-osmolar solutions composed of 1×TE + 100 mM KCl + 87 mM sucrose. DNA-LUV samples, supplemented with the correction buffer, were pipetted into 384 well-plates, covered with MicroAmp Optical Adhesive Film (ThermoFisher Scientific) to prevent evaporation, and allowed to equilibrate and thermalise ($T = 25$°C) inside the microplate reader for at least an hour. Measurements were performed using $\lambda_{\text{excitation}} = 388 \pm 15$ nm and $\lambda_{\text{emission}} = 670 \pm 15$ nm on the "Enhanced Dynamic Range" mode of the instrument, which adjusted the focal height in each independent repeat. In each measurement, a cycle was defined as 20 flashes per well. DNA-LUV samples were measured for 50 cycles, followed by a brief pause for the addition of NMM + G-rich strand. Samples were then measured for 950 further cycles, thereby monitoring NMM fluorescence for a total of $\sim$ 2000 seconds ($\sim$ 30 minutes). In each independent repeat, a control background sample was also monitored, where NMM was added (but omiting the G-rich strand) to a buffered saline solution supplemented with sugar to a composition equal to that of DNA-LUV samples. Fluorescent profiles for each repeat were then processed individually, undergoing background subtraction and normalisation. Averaged fluorescent profiles monitoring the kinetics of assembly of membrane-bound $(G)_5$- and $(G)_6$-nanostructures are presented in Figs. S15 and S16, respectively.

##### Monitoring resorufin production

Stoichiometrically-adjusted hemin co-factor (8.35 $\mu$L), mixed with 10 $\mu$L of our second correction buffer (resulting in osmotically-balanced solutions containing either 1×TE + 87 mM of sugar (sucrose for LUVs and glucose for GUVs) + 100 mM of monovalent salt or 1×TE + 281 mM of sugar + 2 mM divalent salt, as required), was gently added to DNA-decorated LUVs bearing G-rich strands (prepared as detailed in "Membrane Functionalisation with

DNA nanostructures"). Samples were incubated for at least 30 minutes, and then pipetted into 384 well-plates, covered with MicroAmp Optical Adhesive Film (ThermoFisher Scientific) to prevent evaporation and allowed to equilibrate and thermalise ($T = 25°C$) inside the microplate reader for at least an hour. Measurements were performed using $\lambda_{\mathrm{excitation}} = 530 \pm 12.5\,\mathrm{nm}$ and $\lambda_{\mathrm{emission}} = 590 \pm 17.5\,\mathrm{nm}$ on the "Enhanced Dynamic Range" mode of the instrument, which adjusted the focal height in each independent repeat. In each measurement, a cycle was defined as 20 flashes per well. DNA-LUV samples were initially measured for 20 cycles, followed by a brief pause for the addition of AmplexRed (AR) to a final concentration of $\sim 8.5\,\mu$M. Samples were then measured for 30 further cycles, paused for the addition of $H_2O_2$ (final concentration of 0.004%) to trigger peroxidation, and then measured for 950 cycles, thereby monitoring resorufin production for a total of 32035 seconds ($\sim$ 9 hours). Note that both the addition of AR and $H_2O_2$ were done while preserving ionic strength and osmolarity of the samples, and thus all peroxidation reactions proceeded in osmotically-balanced solutions containing either 1×TE + 87 mM of sugar (sucrose for LUVs and glucose for GUVs) + 100 mM of monovalent salt or 1×TE + 281 mM of sugar + 2 mM divalent salt, as required.

Fluorescent profiles, such as those shown in Figs. 3b (main text), S17, S18, and S19, were processed to compute $k_{\mathrm{t_0}}$. To that end, the first 30 data points following the addition of $H_2O_2$ were subjected to linear fitting, from which the slope ($k_{\mathrm{t_0}}$) was extracted. The fold change in conversion rates observed with membrane-bound DNA receptors ($k_{\mathrm{t0,DNAzyme}}$) was quantified relative to the peroxidation rates of hemin co-factor $k_{\mathrm{t0,Hemin}}$ in each of the buffered saline solutions with added sucrose using

$$k^*_{\mathrm{t_0}} = \frac{k_{\mathrm{t0,DNAzyme}}}{k_{\mathrm{t0,Hemin}}} \tag{S3}$$

and construct the heatmaps in Figs. 3c (main text) and S20.

Finally, profiles collected to monitor the activity of DNA-decorated synthetic cells (Fig. 4e) were obtained by normalising fluorescent traces from individual repeats (in Figs. S24 and S25). The profiles were normalised to the intensity values of the first data-point collected after $H_2O_2$ addition. Given the emission/excitation overlap of TexasRed in GUV membranes with resorufin, normalisation allowed to account for the slight differences in GUVs present at the measured well-positions and focal heights, which emerged as a consequence of sample preparation (from batch-to-batch variation and induced by flows from the addition of AmplexRed and hydrogen peroxide).

#### Fluorescence Microscopy of GUVs

DNA-functionalised GUVs were placed in polystyrene 96 well-plates (Greiner Bio-One) and allowed to sediment on the bottom for at least 10 minutes prior to imaging. The chambers were sealed with sticky-tape to prevent evaporation.

##### Confocal Microscopy

Confocal images were acquired at 1400 Hz, averaging over 10 frames, with a Leica TCS SP5 confocal microscope and an HC PL APO CORR CS 40× / 0.85 dry objective from Leica. Alexa488 (excitation maximum - 495 nm; emission maximum - 520 nm) signal was acquired exciting with an Ar-ion laser (488 nm).

##### Epifluorescence Microscopy

Epifluorescence micrographs were acquired after DNAzyme-induced peroxidation to assess the structural stability of our synthetic cell models. GUV-containing well plates were placed on the microscope stage, and the vesicles were allowed to equilibrate for at least $\sim 10$ minutes. Micrographs were acquired using a Nikon Eclipse Ti2-E inverted microscope, equipped with a digital camera (Hamamatsu ORCA-Flash4.0 V3), a tuneable light source (Lumencor

SPECTRA X LED engine), and a Plan Fluor 40× 0.95 N.A. dry objective (Nikon).

### Supplementary Note I: Thermodynamic description of G4 equilibrium assembly

Throughout this section, we describe the equilibrium assembly of tetramolecular G-quadruplexes. We consider individual DNA ($A$) monomers that associate to form G-quadruplexes ($G$).

$$4\,A \rightleftharpoons G$$

The equilibrium expression yields

$$K_{\mathrm{eq}} = \frac{[G]}{[A]^4} = \rho_\circ^{-3}\exp\left(-\frac{\Delta G^\circ}{k_{\mathrm{B}}T}\right) \tag{S4}$$

where $\rho_\circ^3 = 1\,\mathrm{M}$ is a standard reference concentration and $\Delta G^\circ$ is the standard free energy of assembly.

Defining $p$ as the fraction of DNA in G-quadruplexes and $(1-p)$ as the fraction of free monomeric DNA, we obtain for the total DNA concentration ($C$) that $[A] = (1-p)C$ and, accounting for G-quadruplex tetramolecularity, $[G] = (p/4)C$.

Substituting into the equilibrium expression and rearranging, these lead to

$$\frac{4(1-p)^4C^3}{p} = \rho_\circ^3\exp\left(\frac{\Delta G^\circ}{k_{\mathrm{B}}T}\right) \tag{S5}$$

Eq. S5 relates the total concentration of DNA ($C$) to the probability of G-quadruplex formation ($p$) and the standard free energy of assembly $\Delta G^\circ$. Finally, we compute numerical solutions for $p$ with $0 < p < 1$ for a range of total DNA concentrations. These are summarised in Fig. S14

#### Supplementary Note II: Estimation of DNA nanostructure local concentration on membrane surfaces

In this section, we derive an estimate of the local concentration ($C_{\mathrm{local}}$) of DNA nanostructures when tethered to lipid membranes using

$$C_{\mathrm{local}} = \frac{n_{\mathrm{DNA}}}{V} \tag{S6}$$

where $V$ is the possible volume that $n_{\mathrm{DNA}}$ moles of membrane-bound DNA nanostructures can occupy.

We consider $V$ to be the available bilayer surface area ($A_{\mathrm{bilayer}}$) multiplied by the length of the 56-bp DNA nano-devices ($L \approx 19\,\mathrm{nm}$). In turn, to determine $A_{\mathrm{bilayer}}$, we use an average area per DOPC lipid of $a = 0.72\,\mathrm{nm}^2$ [11] and assume no lipid loss throughout LUV preparation (that is, all lipids self-assemble into bilayer membranes, of which only the outer leaflet is accessible for functionalisation). In this manner, at the relevant volumes and concentrations of LUVs ($9.2\,\mu$L at $C = 4\,\mathrm{mg\ mL}^{-1}$) and DNA nanostructures ($16.7\,\mu$L at $C = 2\,\mu$M) in our experimental implementation, we find

$$C_{\mathrm{local}} = \frac{3.34 \times 10^{-11}\mathrm{moles}}{1.95 \times 10^{17}\mathrm{nm}^3} = 171.7\,\mu\mathrm{M} \tag{S7}$$

#### Supplementary Note III: Confocal micrographs acquisition and analysis

Similar to our previous contributions,[5,6] confocal micrographs were processed to assess the domain-accumulation tendencies of our DNA nano-devices. We thus acquired micrographs of fields of view of $64.84\,\mu m \times 64.84\,\mu m$ in size. Z-stacks for 3D reconstructions, in Figs. 4 (main text) and S15, were acquired with a slice thicknesses of $0.5\,\mu m$. The size of the field of view is the largest our instrument can image at the fastest scanning rate of 1400 Hz. Screening and selection of imaged vesicles was underpinned by two criteria:

1. Vesicles were smaller than the size of the field of view and big enough such that Brownian motion was negligible within the acquisition timescales of a few minutes per GUV.

2. The orientation of the vesicle was such that a sharp interface separating the $L_\mathrm{d}$ and $L_\mathrm{o}$ domains was visible at the equatorial cross-section of the GUV, thus enabling quantitative processing.

#### Data Analysis

An image segmentation pipeline, developed in a previous contribution,[5,12] was applied to process equatorial micrographs of DNA-decorated GUVs and quantify the thermodynamic tendencies of $L_\mathrm{o}$-partitioning for DNA nanostructures tethered to membranes through double-cholesterol (dC) anchors. Briefly, the membrane contour was segmented following a noise removal and line profiling routine, allowing to sample the signal of fluorescein-labelled nanostructures. Average fluorescence intensity values along the membrane circumference were determined through Gaussian fitting and numerical integration, and subsequently classified as belonging to $L_\mathrm{o}$ or $L_\mathrm{d}$ phases by comparing their location to that of the TexasRed-DHPE signal, a fluorescent lipid marker which preferentially stains $L_\mathrm{d}$-domains. The $L_\mathrm{o}$ and $L_\mathrm{d}$-associated DNA intensities ($I_{L_\mathrm{o}}$ and $I_{L_\mathrm{d}}$, respectively), were used to compute a fractional intensity using

$$f_{\mathrm{p},L_\mathrm{o}} = \frac{I_{L_\mathrm{o}}}{(I_{L_\mathrm{o}} + I_{L_\mathrm{d}})} \tag{S8}$$

which describes the lateral organisation of DNA nanostructures in the membrane.

To better quantify the effect of receptor assembly on lateral distribution, we computed the fold enhancement in $L_{\mathrm{o}}$-partitioning coefficients ($K_{\mathrm{p}} = I_{L_{\mathrm{o}}}/I_{L_{\mathrm{d}}}$) of $(\mathrm{G})_6$-decorated membranes relative to GUVs functionalised with DNA nano-devices lacking the G-rich strand using

$$K_{\mathrm{p}}^{*} = \frac{K_{\mathrm{p,(G)_6-DNA-GUVs}}}{K_{\mathrm{p,DNA-GUVs}}} \tag{S9}$$

which we plot in Fig. 4c (main text) for GUVs functionalised in monovalent ($\mathrm{K}^{+}$, $\mathrm{Na}^{+}$, or $\mathrm{Li}^{+}$) or divalent ($\mathrm{Ca}^{2+}$ or $\mathrm{Mg}^{2+}$) cationic conditions. Finally, we applied a non-parametric, one-tailed, Wilcoxon-signed rank test and determined that the resulting $K_{\mathrm{p}}^{*}$ distributions were significantly higher than 1 across all cationic conditions (statistical significance with $p < 1.2 \times 10^{-16}$, see $p$-values in Table S4).

### Supplementary Figures

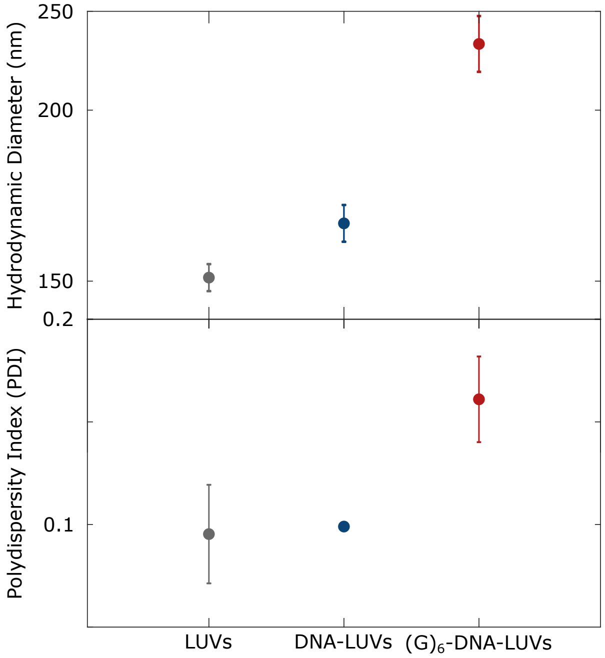


Figure S1: **LUV hydrodynamic sizes upon membrane functionalisation suggest a low degree of inter-vesicle linking mediated by G-quadruplexes.** Mean ± standard deviation of hydrodynamic diameter (top) and polydispersity indices (PDIs, bottom) of LUV populations as obtained with DLS measurements at the various stages of the membrane functionalisation process: LUVs, DNA-LUVs, and $(G)_6$-DNA-LUVs; see Figure 1b for Intensity-based distributions. The moderately higher PDI values for $(G)_6$-DNA-LUVs, consistent with a slight broadening in size distribution, suggest that – while LUV dispersions are predominantly monodisperse – there is a low degree of inter-vesicle association facilitated by G-quadruplexes. Given the limited broadening of the size distribution and the sample monodispersity, we conclude that intra-vesicle receptor assembly is the dominant mode of interactions, with only minor inter-vesicle assemblies.

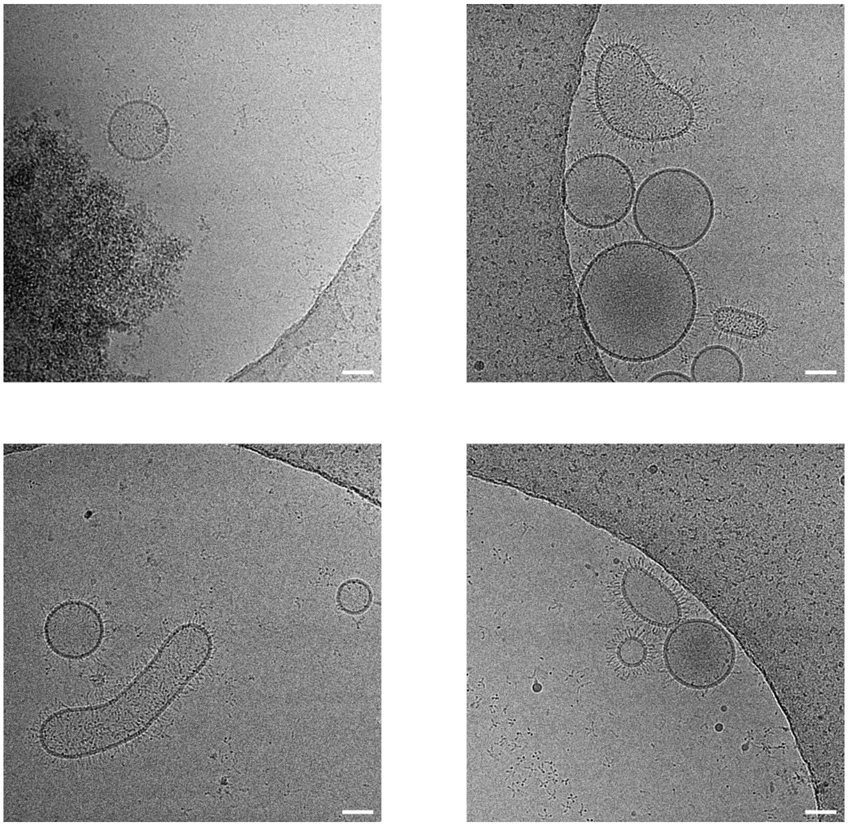

Figure S2: **Cryo-electron micrographs of membranes decorated with G4-forming DNA nano-devices.** Representative cryo-electron micrographs of LUVs decorated with DNA nano-devices carrying $(G)_6$-overhangs, where G-quadruplexes manifest as membrane-localised electron-dense regions. We observe background signal from DNA constructs that are not membrane-bound, which we ascribe to bilayer disruption and DNA nanostructure detachment during sample preparation, particularly due to the mechanical shear forces involved in blotting.[13] Given the double-cholesterol anchors, we do not expect thermal unbinding of the nano-devices, as each anchor contributes a free energy of insertion in the order of $\approx 20$ to $30\,k_\mathrm{B}T$,[14,15] rendering membrane tethering effectively irreversible at room temperature. Scale bars = 50 nm.

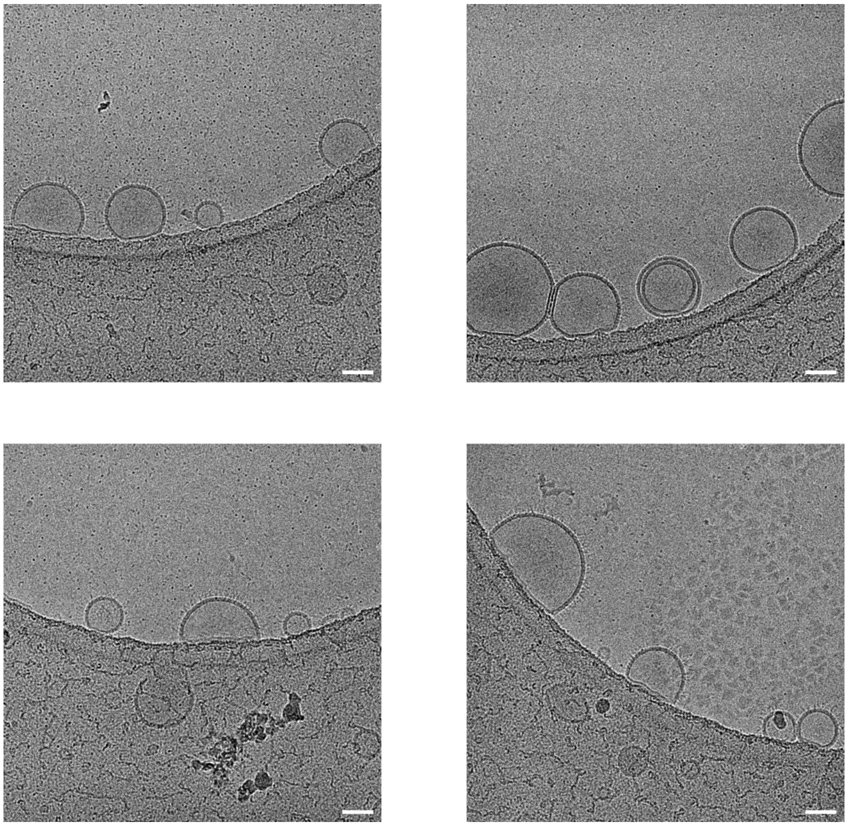

Figure S3: **Cryo-electron micrographs of membranes decorated with DNA nano-devices.** Representative cryo-electron micrographs of LUVs decorated with DNA nano-devices that lack the $(G)_6$-strand, resulting in shorter nano-devices and the absence of membrane-localised electron-dense regions. We observe background signal from DNA constructs that are not membrane-bound, which we ascribe to bilayer disruption and DNA nanostructure detachment during sample preparation, particularly due to the mechanical shear forces involved in blotting.[13] Given the double-cholesterol anchors, we do not expect thermal unbinding of the nano-devices, as each anchor contributes a free energy of insertion in the order of $\approx 20$ to $30\,k_\mathrm{B}T$,[14,15] rendering membrane tethering effectively irreversible at room temperature. Scale bars = 50 nm.

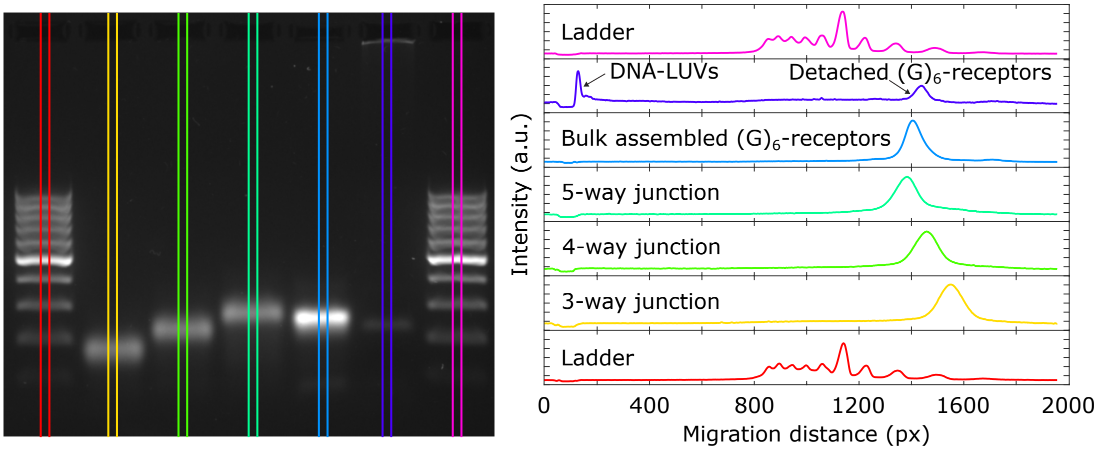


Figure S4: **Gel-shift assay confirms tetrameric stoichiometry of biomimetic DNA receptors.** Comparing the electrophoretic mobilities of multivalent DNA junctions with that of G4-stabilised nanostructures, either assembled in the bulk via thermal annealing or on the membrane surface and subsequently detached through a strand displacement reaction,[16] confirms stoichiometry of assembly. The similar migration profile observed for our G4-nanostructures to that of a 4-way junction with arms of equal size to that of monomeric DNA constructs indicates that receptor stoichiometry is tetrameric. The slight reduction in receptor mobility relative to 4-way junctions likely emerges due to the flexible linker between G-runs and membrane-anchoring duplexes, which is absent from the central junction of the control multivalent nanostructures.

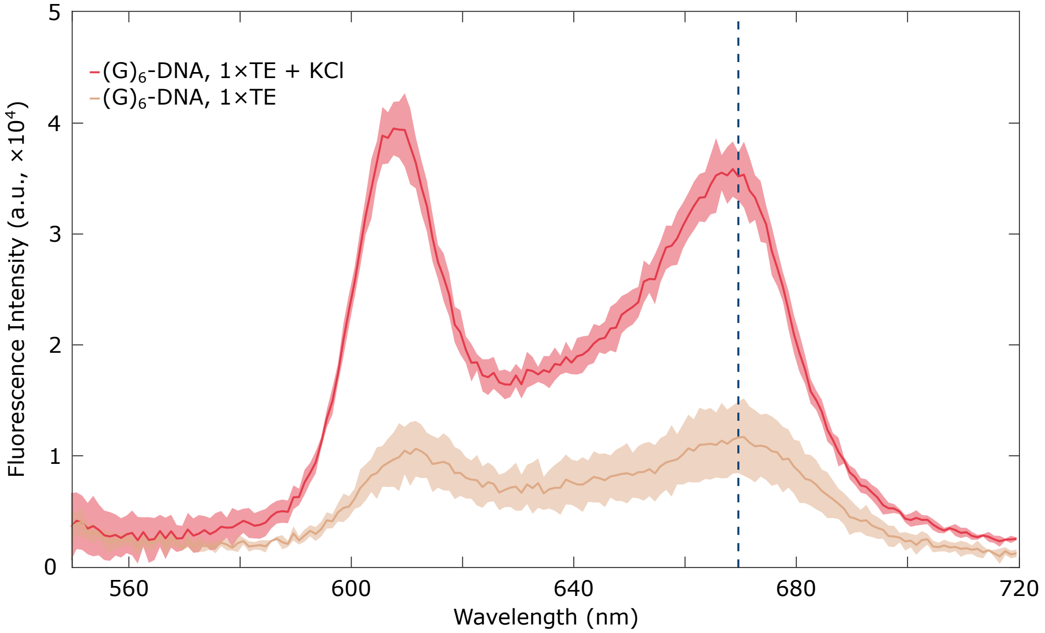


Figure S5: **The fluoresence intensity of NMM increases specifically in the presence of G-quadruplexes.** Fluorescence emission spectra of NMM in the presence of $(G)_6$-DNA nanostructures dispersed in 1×TE buffering solutions either supplemented with or lacking 100 mM KCl. When in the presence of $K^+$ ions, G-quadruplex formation results in an enhancement of NMM fluorescence intensity. Note that no spectral peak shift is observed, particularly at the wavelength used to monitor NMM fluorescence ($\lambda = 670$ nm, marked by the blue dashed line).

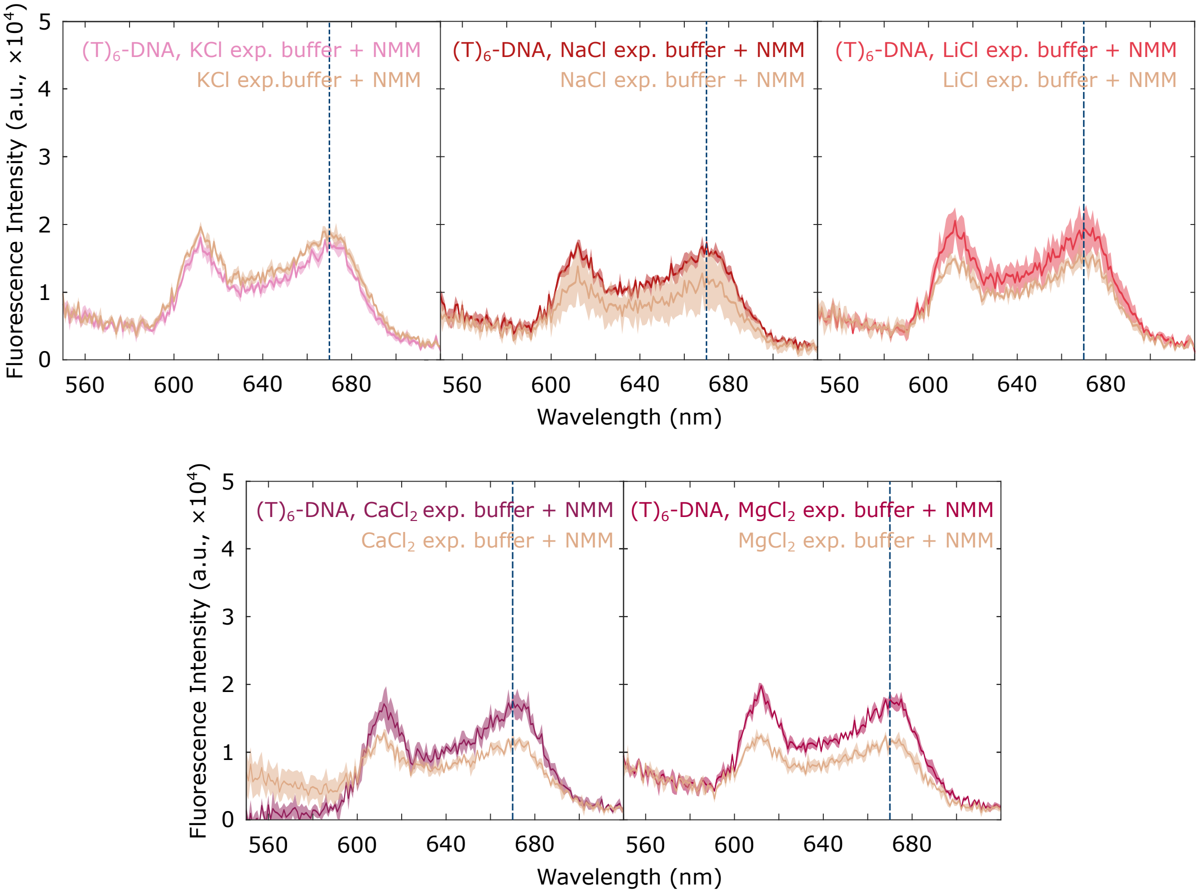


Figure S6: **NMM fluorescence emission spectrum does not shift with cationic composition in the presence of DNA nanostructures.** Fluorescence emission spectra of NMM dispersed in our experimental buffer (i.e. either 1×TE + 100 mM monovalent salt + 87 mM sucrose or 1×TE + 2 mM divalent salt + 281 mM sucrose) in the presence or absence of $(T)_6$-DNA nanostructures, which cannot assemble into G-quadruplexes. While the presence of DNA nanostructures leads to modest increase in NMM fluorescence intensity (as also shown in Fig. S5), no spectral shifts are observed and our choice of wavelength ($\lambda = 670\,$nm, marked by the blue dashed line) remains appropriate.

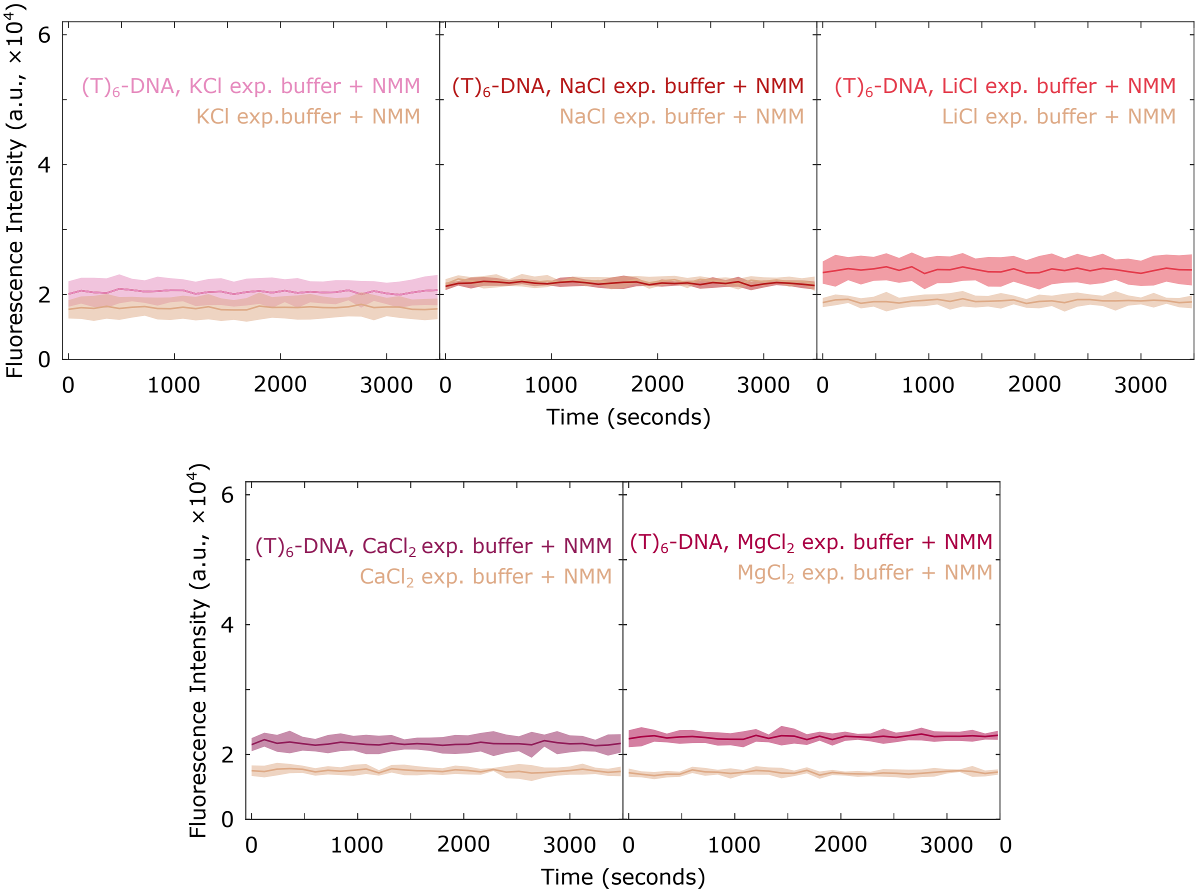


Figure S7: **NMM fluorescence intensity has a minor dependence on cationic composition.** Fluorescence intensity of NMM dispersed in our experimental buffer (i.e. either $1\times$TE + 100 mM monovalent salt + 87 mM sucrose or $1\times$TE + 2 mM divalent salt + 281 mM sucrose) in the presence or absence of (T)$_6$-DNA nanostructures. The presence of these DNA nanostructures, that do not form G-quadruplexes, results in modest changes in NMM fluorescence intensity, which depend on cationic composition.

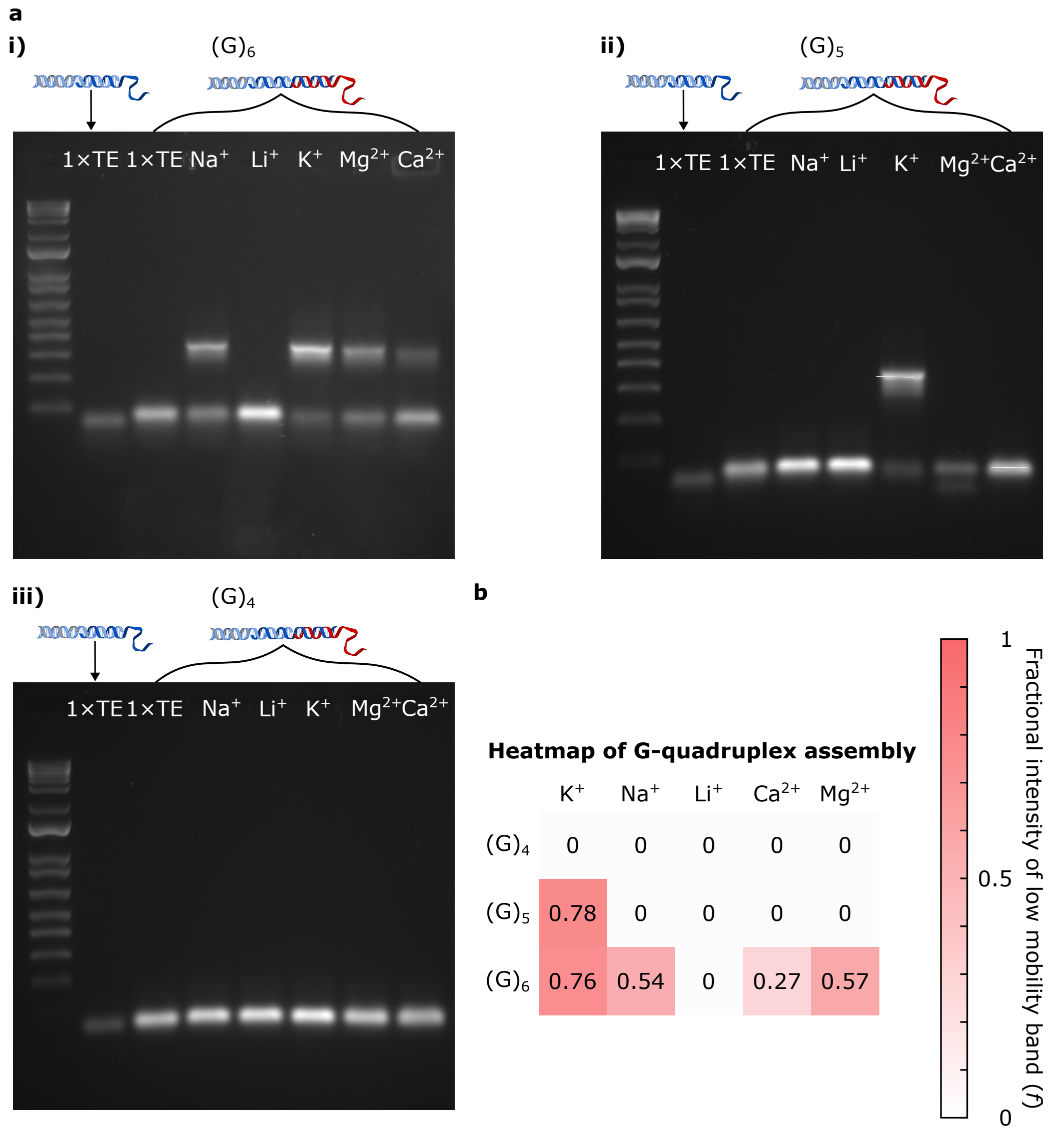


Figure S8: **G-quadruplex formation quantified with AGE for various cationic conditions. a** Micrographs of agarose gel electrophoresis of non-cholesterolised nano-devices lacking or featuring (G)-rich strands (schematically depicted in red) and dispersed in buffered solutions: either 1×TE or 1×TE with added monovalent (NaCl, KCl, or LiCl at [Salt] = 100 mM) or divalent ($MgCl_2$ or $CaCl_2$ at [Salt] = 2 mM) cations. **b** Heatmap of fractional intensity of the low-mobility band ($f$), computed as detailed in the Experimental Section ("Agarose Gel Electrophoresis"). The heatmap replicates the assembly trends summarised in Fig. S4. Note that in the $Mg^{2+}$ lane (panel a-ii), the second band, below that expected for monomeric G-rich nanostructures, possibly emerged due to imbalances in the stoichiometry between G-rich strand and the strands that compose the (non-cholesterolised) anchor device (in blue).

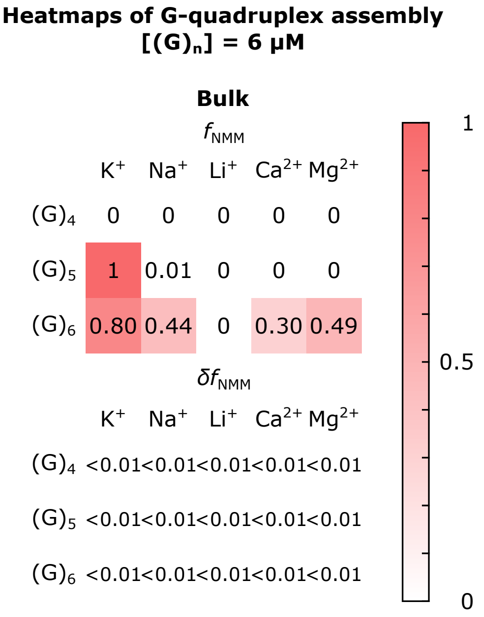


Figure S9: **Bulk self-assembly of G-quadruplexes assessed with NMM fluorimetry.** Heatmaps of fractional NMM intensity ($f_{\mathrm{NMM}}$, Top) and its standard deviation ($\delta f_{\mathrm{NMM}}$, Bottom), computed as detailed in the Experimental Section ("Monitoring the degree of tetramer assembly with NMM fluorimetry") exploring designs $(\mathrm{G})_{\mathrm{n}=4,5,6}$ and cationic conditions for non-cholesterolised nanostructures dispersed in the bulk at concentrations that favour G-quadruplex formation $[(\mathrm{G})_{\mathrm{n}}] = 6\,\mu\mathrm{M}$ in buffered saline solutions ($1\times$TE + 100 mM monovalent salt or $1\times$TE + 2 mM divalent salt).

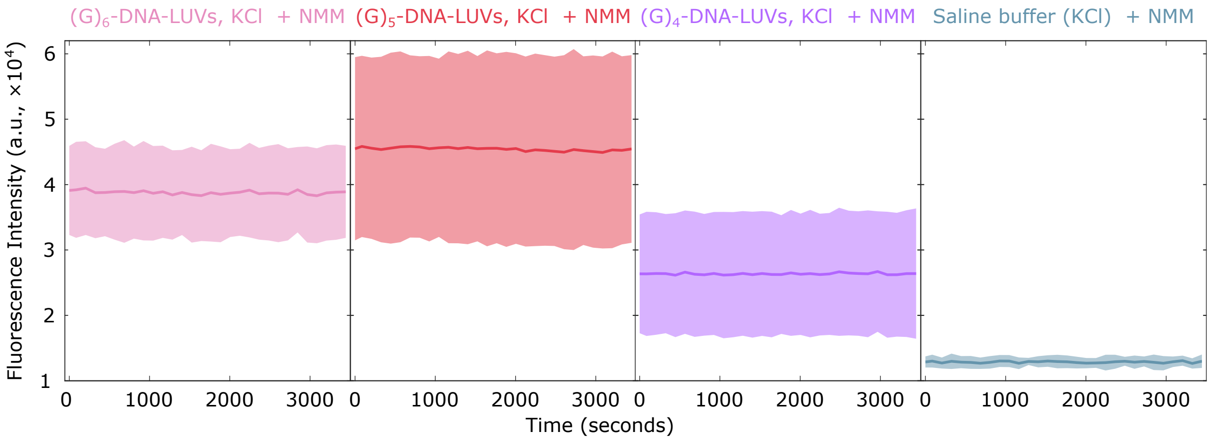


Figure S10: **Representative NMM fluorescence profiles show equilibrium binding.** Time-invariant fluorescent profiles of DNA-decorated LUVs in various cationic conditions: $(G)_{n=4,5,6}$ in KCl, relative to the background signal of NMM dispersed in saline buffer supplemented with sugar (1×TE + 100 mM KCl + 87 mm sucrose). Fluorescent profiles show no intensity changes within timescales of our measurements ($\sim$ 1 hour), thus indicating that NMM binding to G-tetrads, as well as G-quadruplex formation, has reached equilibrium.

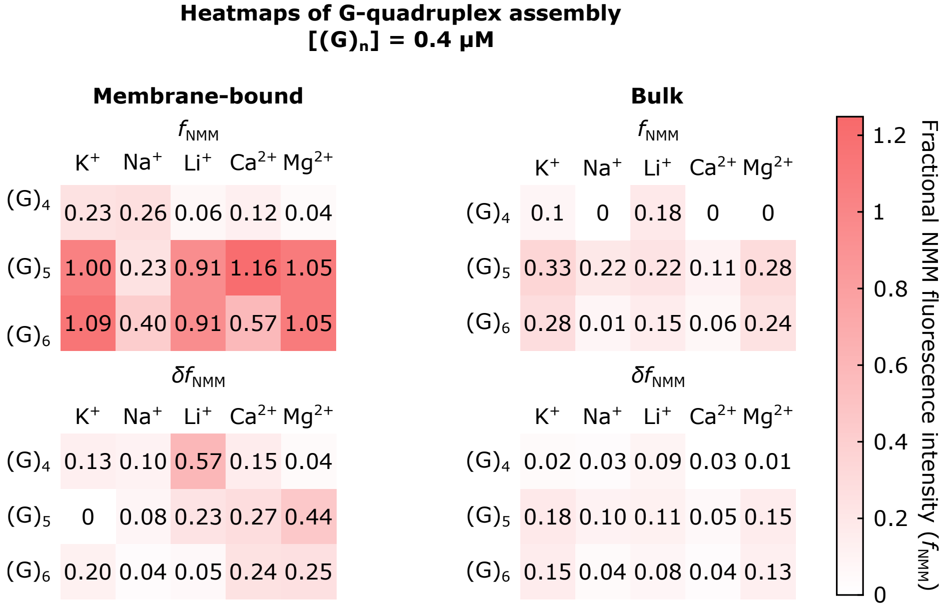


Figure S11: **G-quadruplex formation is enhanced when membrane-confined relative to assembly in the bulk.** Heatmaps of fractional NMM intensity ($f_{\mathrm{NMM}}$, Top) and its standard deviation ($\delta f_{\mathrm{NMM}}$, Bottom), computed as detailed in the Experimental Section ("Monitoring the degree of tetramer assembly with NMM fluorimetry") exploring designs $(\mathrm{G})_{\mathrm{n}=4,5,6}$ and various cationic conditions for: membrane-bound DNA nanostructures (prepared as described in the Experimental Section "Membrane Functionalisation with DNA nanostructures); and for non-cholesterolised nanostructures dispersed in the bulk at nominally equal concentrations $[(\mathrm{G})_{\mathrm{n}}] = 0.4\,\mu\mathrm{M}$ in buffered saline solutions supplemented with sucrose and lacking LUVs. Note that for the bulk scenario, sample preparation followed a sequential implementation similar to that of DNA-LUVs, where non-cholesterolised nanostructures were first dispersed in buffered saline solutions with sucrose, and subsequently G-strands were added to allow for hybridisation. Samples were incubated overnight before the fluorimetry measurements.

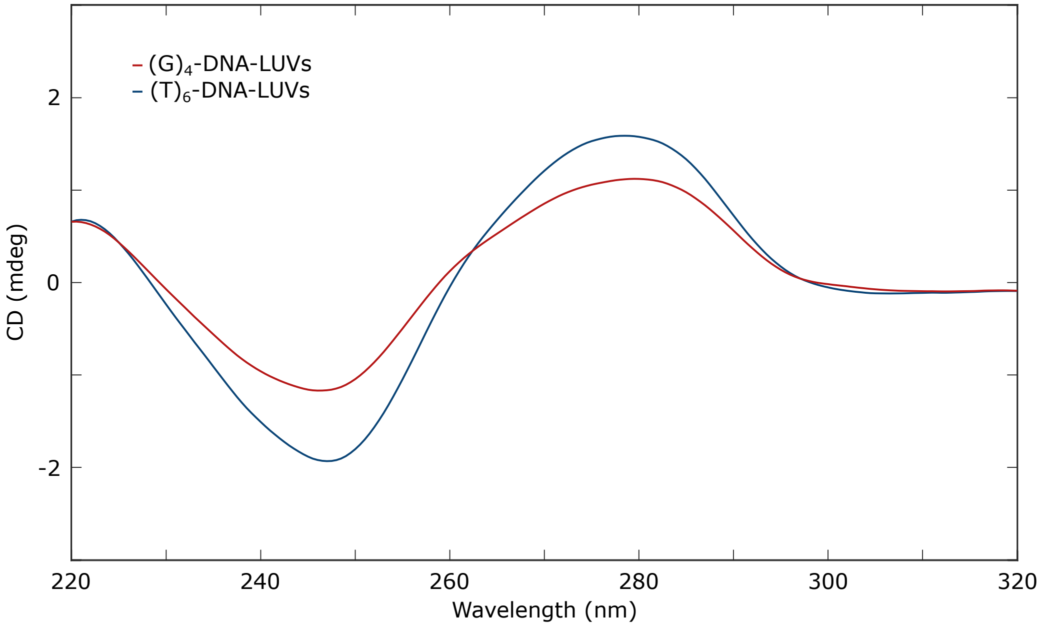


Figure S12: **Circular Dichroism of (G)$_4$-functionalised LUVs suggests moderate assembly of parallel G-quadruplexes.** Moderate peak shifting in circular dichroism spectra of (G)$_4$-functionalised LUVs (red) relative to LUVs functionalised with DNA nanostructures, but lacking the G-rich overhang (blue), suggests a moderate tendency of (G)$_4$-overhangs to assemble G-quadruplexes on the surface of lipid membranes.

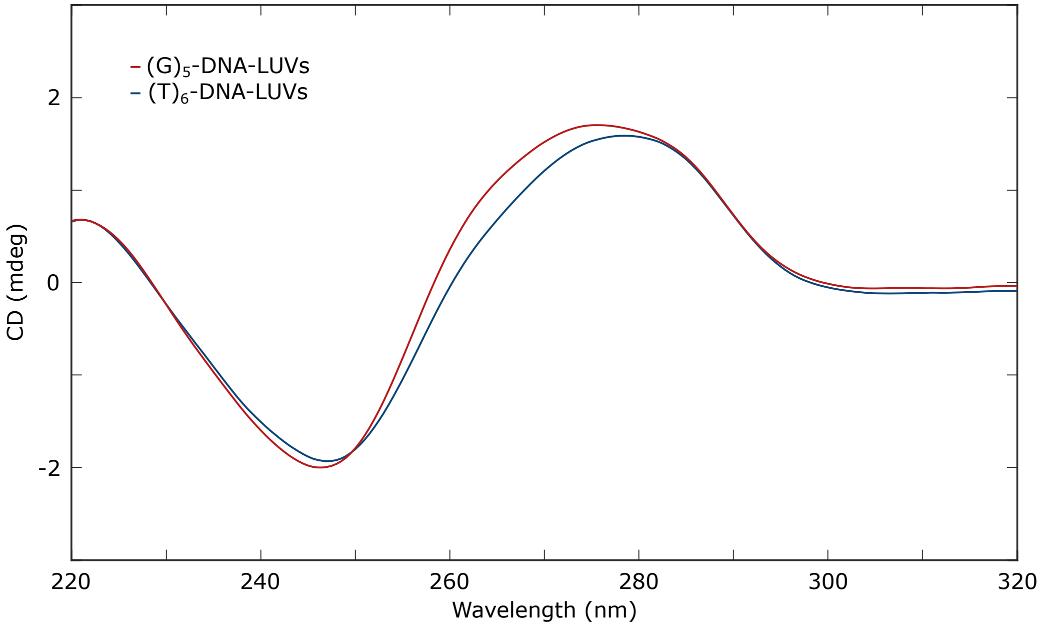


Figure S13: **Circular Dichroism of $(G)_5$-functionalised LUVs indicates assembly of parallel G-quadruplexes.** Characteristic peak shifts in circular dichroism spectra of $(G)_5$-functionalised LUVs (red) relative to LUVs functionalised with DNA nanostructures, but lacking the G-rich overhang (blue), indicate the presence of parallel G-quadruplexes on the surface of lipid membranes.

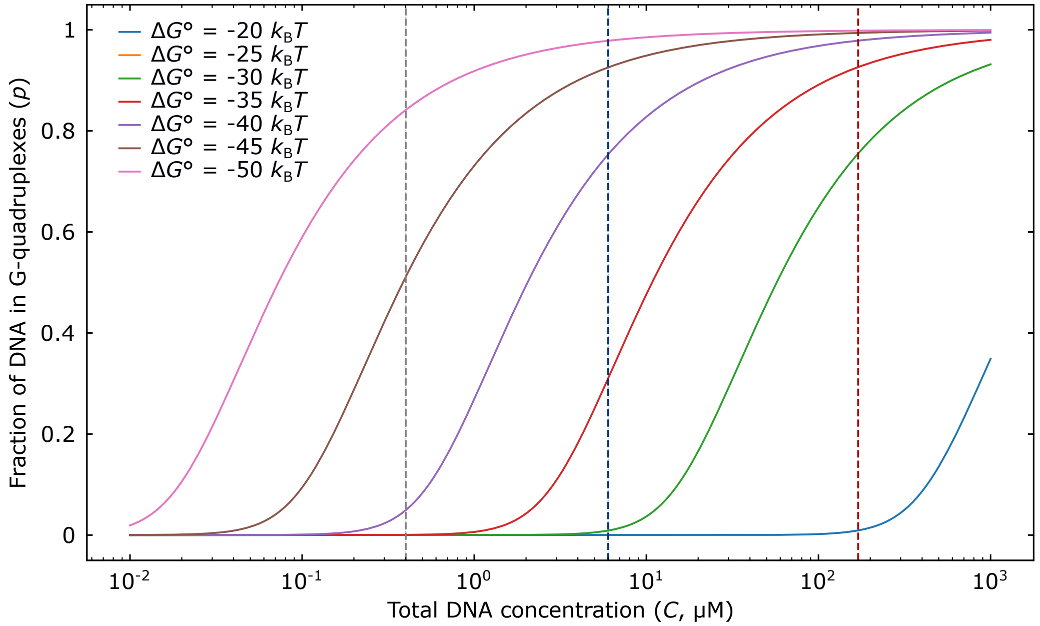


Figure S14: **Equilibrium assembly of tetramolecular G-quadruplexes is highly sensitive to DNA concentration.** Using a simple thermodynamic description of tetrameric assembly (see Supplementary Note I), we relate the fraction of DNA constructs in G-quadruplexes ($p$) to the total concentration of DNA nanostructures ($C$) and the standard free energy of assembly ($\Delta G^{\circ}$). The resulting plot shows how, at specific $\Delta G^{\circ}$ values, nominally low concentrations, such as those in bulk assays (e.g. $0.4\,\mu$M, marked by the dashed gray line or $6\,\mu$M, marked by the dashed blue line), probabilities of G4 formation tend to be lower. In turn, high DNA concentrations, such as those induced by membrane-anchoring ($171\,\mu$M, marked by the dashed red line), correspond to a substantially higher $p$ values. For instance, consider a representative scenario with $\Delta G^{\circ} = -30 k_{\mathrm{B}}T$. At these low DNA concentrations, $p$ is negligible, while under membrane-confinement the local DNA concentration corresponds to $p \sim 0.7$, a 14-fold enhancement in G-quadruplex formation probability.

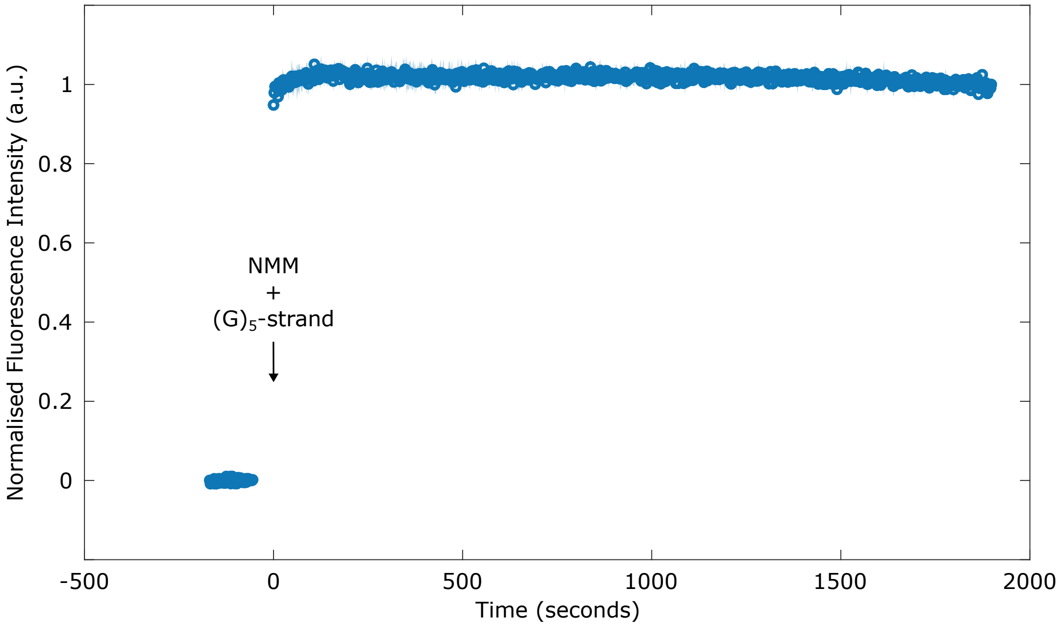


Figure S15: **Assembly kinetics of membrane-bound (G)$_5$-nano-devices.** Normalised NMM fluorescent profile following the addition of (G)$_5$-strands to DNA-decorated LUVs in the presence of KCl to monitor the kinetics of G-quadruplex assembly on the surface of lipid membranes (see Experimental Section). Markers represent the mean, while the shaded region represents the standard deviation, of n = 3 independent replicates. Each independent profile has been background-subtracted (relative to the addition of NMM to saline buffers with added sugar), and averaged after setting $t = 0$ as the first data-point collected after addition. Note that fluorescence values equilibrate in under 250 s, likely limited by the timescales of diffusion of oligonucleotides to bind the DNA anchors on the membrane. In turn, these timescales suggest that G-quadruplex assembly occurs on a much faster timescale, consistent with the high local concentrations of DNA nanostructures at the membrane surface, estimated in Supplementary Note I to be $\sim 170\,\mu$M].

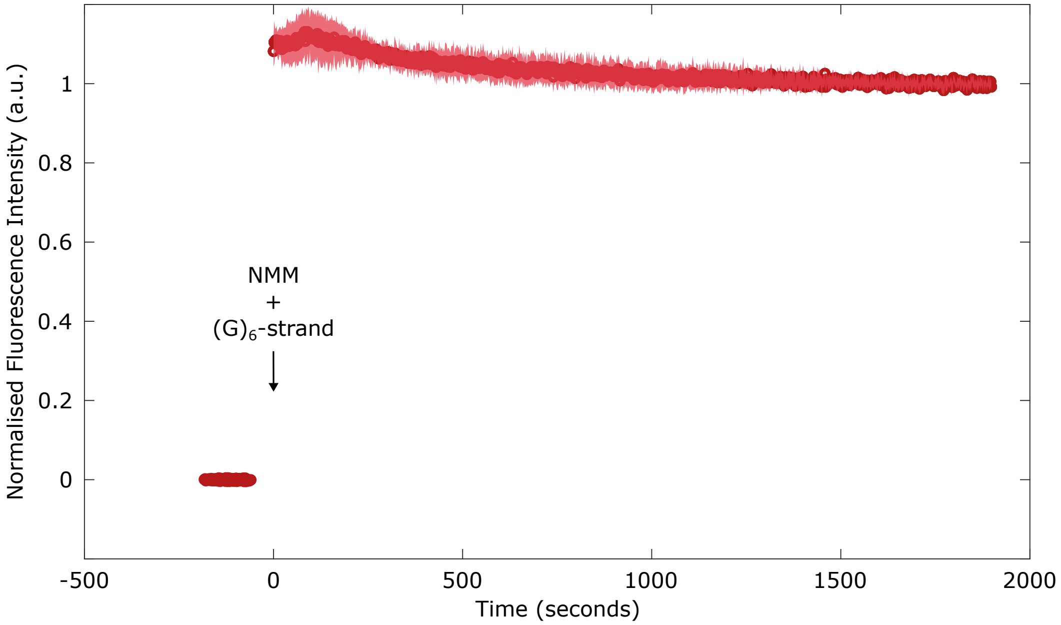


Figure S16: **Assembly kinetics of membrane-bound $(G)_6$-nano-devices.** Normalised NMM fluorescent profile following the addition of $(G)_6$-strands to DNA-decorated LUVs in the presence of KCl to monitor the kinetics of G-quadruplex assembly on the surface of lipid membranes (see Experimental Section). Markers represent the mean, while the shaded region represents the standard deviation, of n = 5 independent replicates. Each independent profile has been background-subtracted (relative to the addition of NMM to saline buffers with added sugar), and averaged after setting $t = 0$ as the first data-point collected after addition. Note that fluorescence values equilibrate in under 250 s, likely limited by the timescales of diffusion of oligonucleotides to bind the DNA anchors on the membrane. In turn, these timescales suggest that G-quadruplex assembly occurs on a much faster timescale, consistent with the high local concentrations of DNA nanostructures at the membrane surface, estimated in Supplementary Note I to be $\sim 170\,\mu$M].

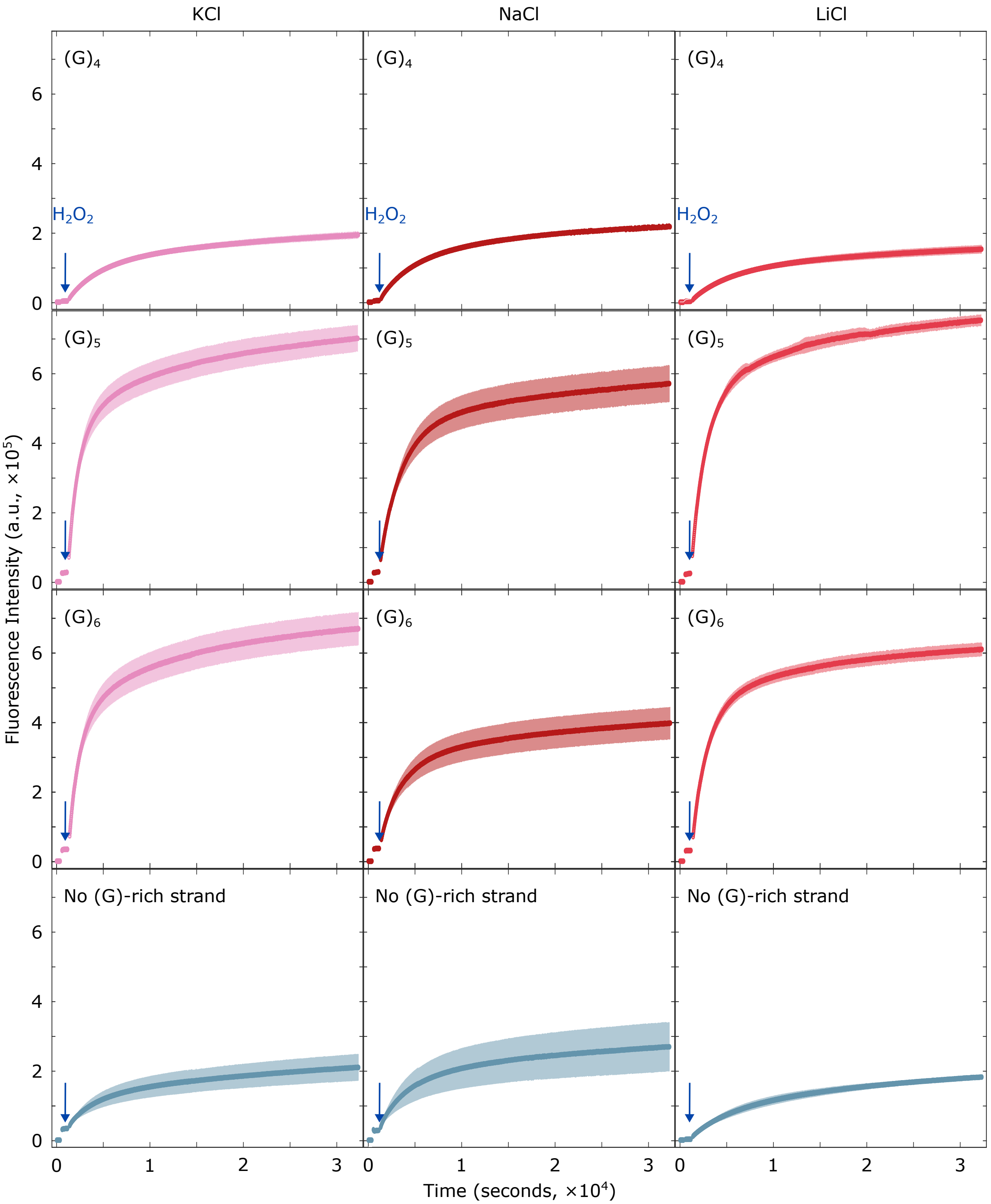


Figure S17: **Rate of resorufin production is modulated with membrane-bound DNA receptors.** Fluorescence intensity profiles of resorufin production when catalysed by membrane-bound hemin/G-quadruplex receptors. Different cationic conditions lead to a wide range of rates, as seen for KCl (left), NaCl (middle) and LiCl (right) for nano-devices featuring design variants $(G)_{n=4,5,6}$ (labelled for each plot). Peroxidation catalysed by DNA receptors is compared to that driven hemin co-factors in the absence of G-rich strands, and therefore, G-quadruplexes. Solid lines represent mean values, while shaded regions are standard deviations, of n = 3 replicates.

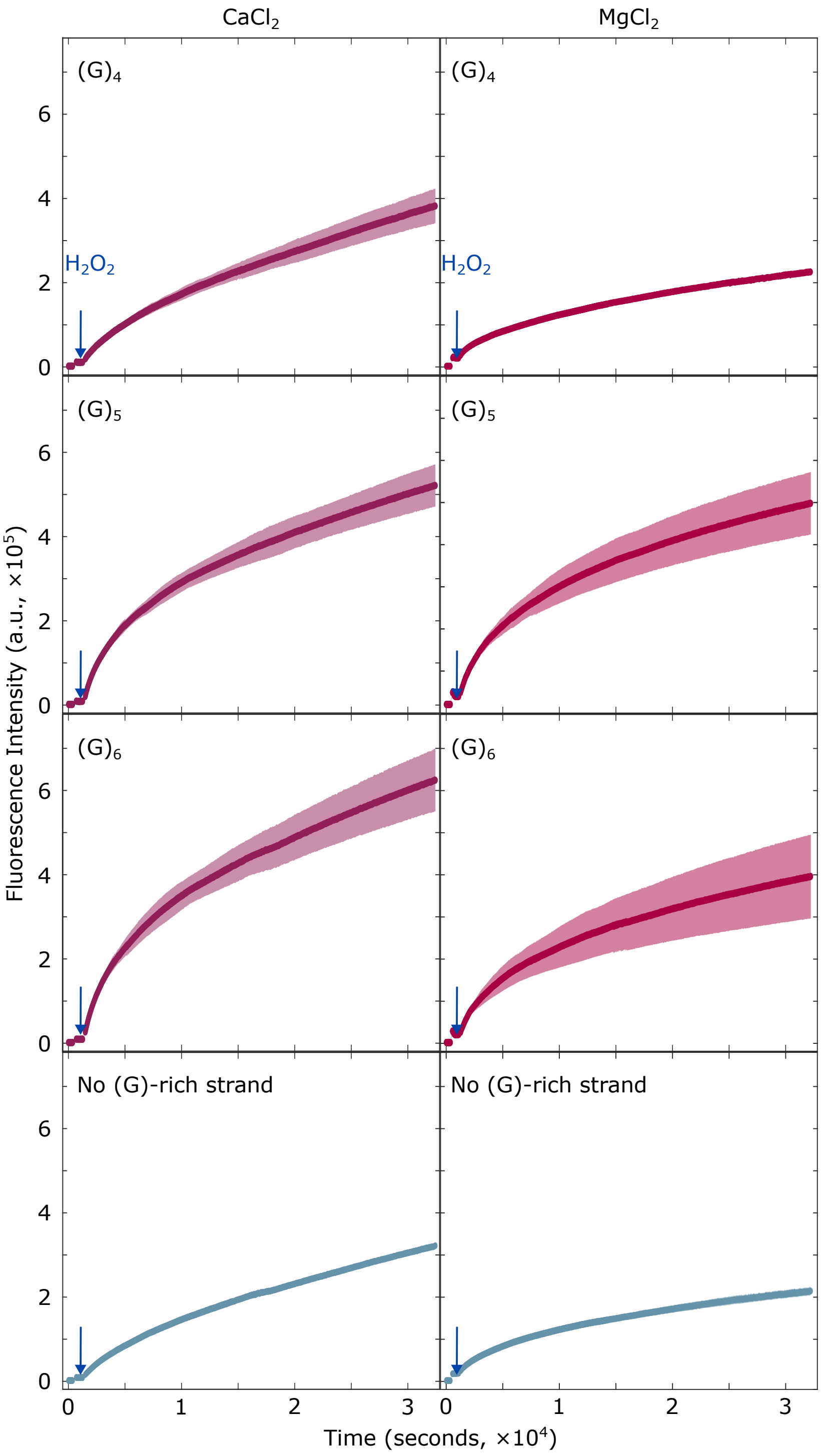


Figure S18: **Rate of resorufin production is modulated with membrane-bound DNA receptors.** Fluorescence intensity profiles of resorufin production when catalysed by membrane-bound hemin/G-quadruplex receptors. Different cationic conditions lead to a wide range of rates, as seen for $CaCl_2$ (left) and $MgCl_2$ (right) for nano-devices featuring design variants $(G)_{n=4,5,6}$ (labelled for each plot). Peroxidation catalysed by DNA receptors is compared to that driven hemin co-factors in the absence of G-rich strands, and therefore, G-quadruplexes. Solid lines represent mean values, while shaded regions are standard deviations, of n = 3 replicates.

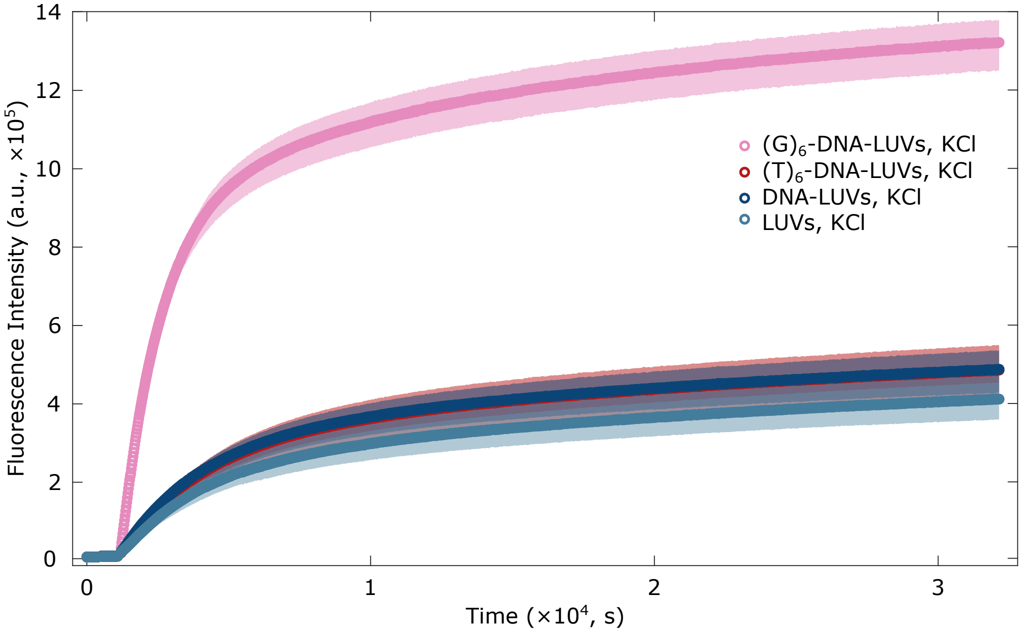


Figure S19: **Hemin peroxidase activity is only enhanced in the presence of G-quadruplexes.** Comparing peroxidation in samples of non-functionalised LUVs as well as LUVs featuring either $(G)_6$-DNA, $(T)_6$-DNA, or simply DNA anchor modules confirms that the enhancement of catalytic activity is a result of G-quadruplex formation rather than non-specific interactions of hemin with unstructured or duplex DNA constructs.

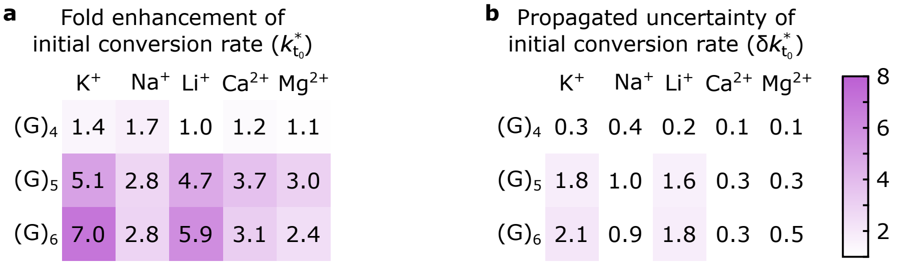


Figure S20: **Fine tuning the assembly of hemin/G-quadruplex DNAzyme receptors affords control over initial peroxidation rates.** Heatmaps of fold change in the initial conversion rate ($k^*_{t_0}$, panel **a**) and its propagated error ($\delta k^*_{t_0}$, **b**), computed as detailed in the Experimental Section ("Monitoring resorufin production") for membrane-bound DNA receptors assembled designs $(G)_{n=4,5,6}$) and exploring various cationic conditions.

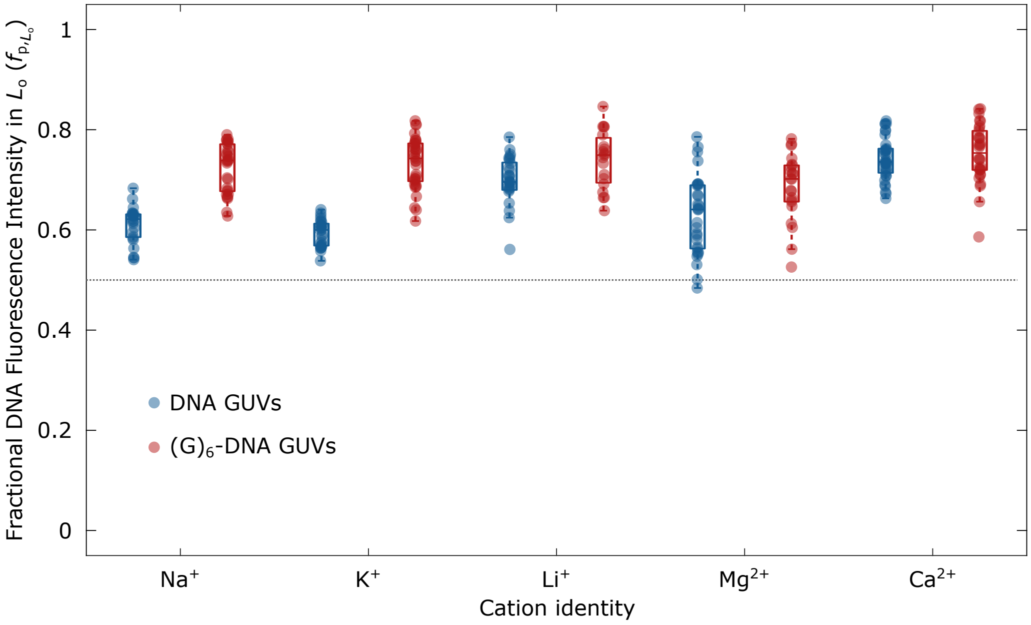


Figure S21: **(G)$_6$-overhangs influence $L_o$-domain partitioning of nano-devices via G-quadruplex formation.** Plots of fractional DNA intensity in $L_o$ ($f_{p,L_o}$), computed as detailed in Supplementary Note I, for GUVs decorated with DNA nanostructures lacking (in blue) or featuring (in red) (G)$_6$-strands as a function of buffered saline solutions (1×TE + NaCl, KCl, LiCl, $MgCl_2$ or $CaCl_2$) with added glucose.

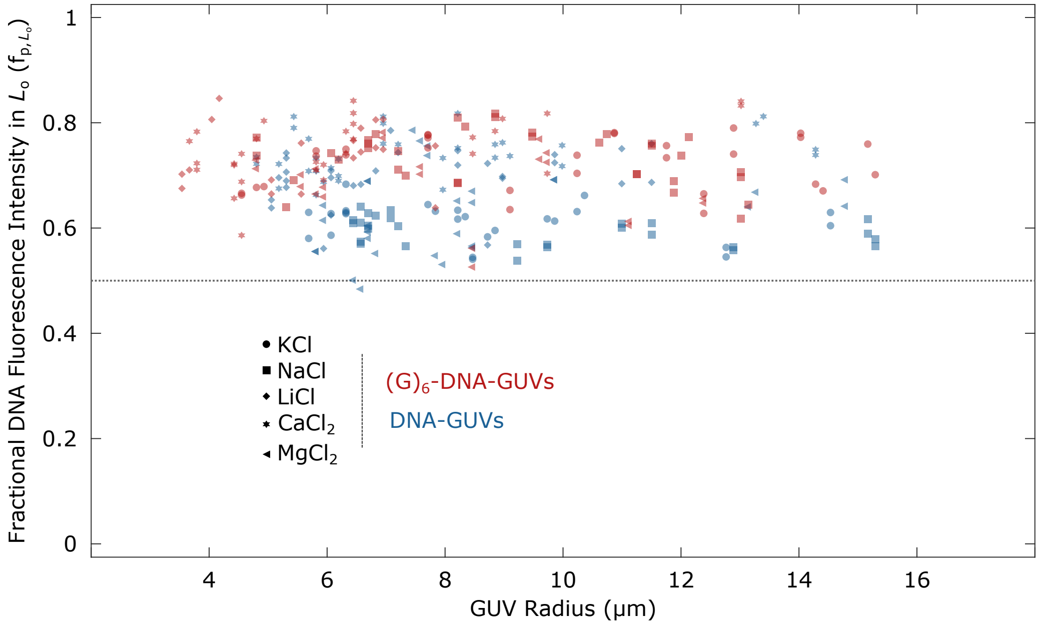


Figure S22: **Vesicle size does not influence accumulation of DNA nanostructures in lipid domains.** Exploring the relationship between the fractional DNA intensity in $L_o$ domains as a function of GUV size revealed that there is no correlation between the partitioning of DNA nanostructures and vesicle size across different cationic compositions (non-parametric Spearman statistical test. See Table S5 for $\rho$ and $p-$values).

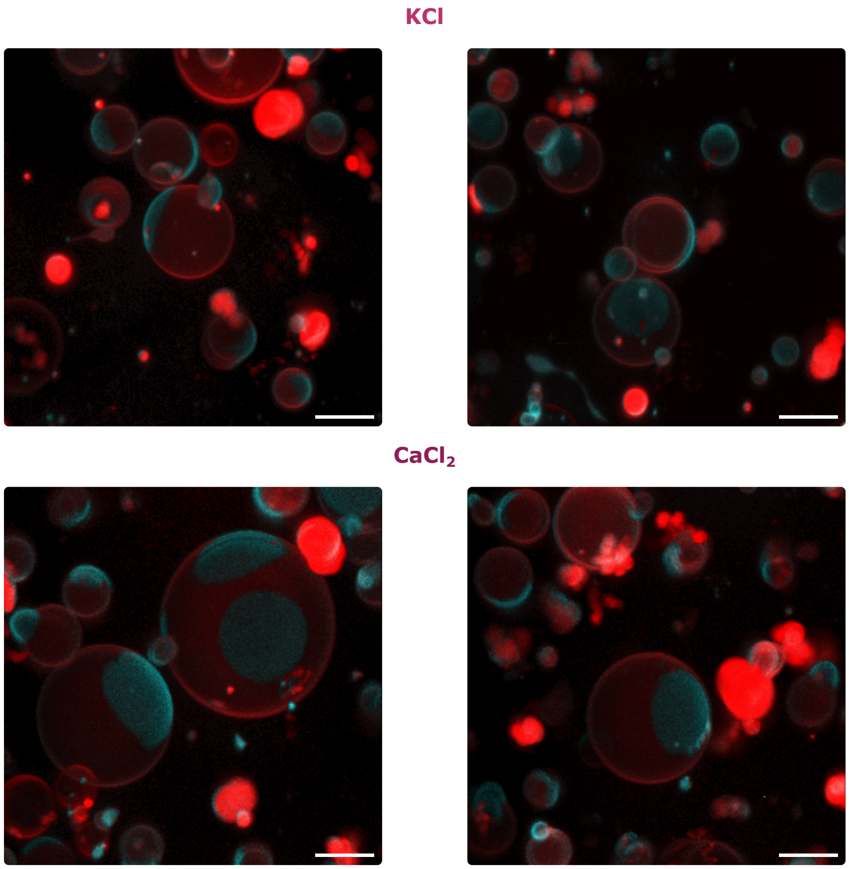


Figure S23: **3D views of synthetic cells decorated with DNA receptors for localised activity.** Representative 3D views (obtained from Volume Viewer, FIJI[17]) reconstructed from confocal z-stacks of synthetic cells decorated with $(G)_6$-DNA receptors in the presence of KCl (top) or $CaCl_2$ (bottom) as monovalent or divalent cation messengers, respectively. Scale bars = 10 $\mu$m.

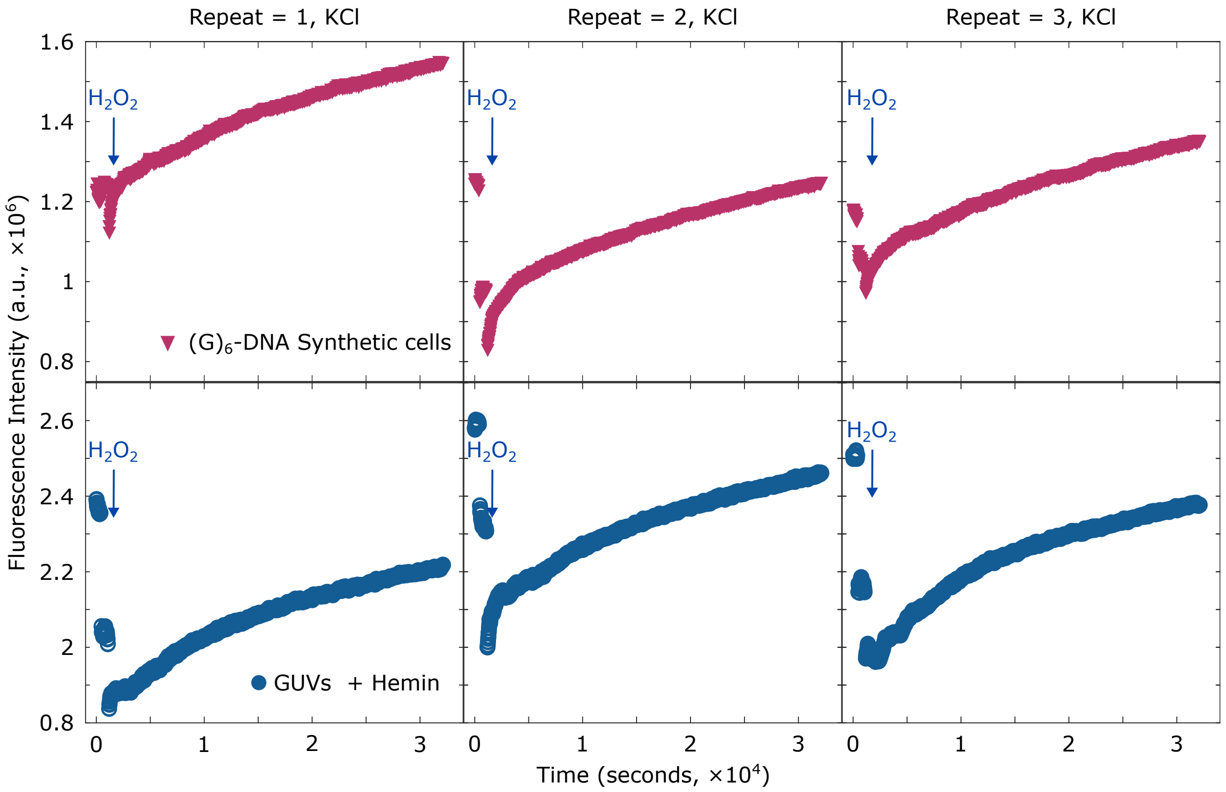


Figure S24: **Peroxidation activity localised to lipid domains of synthetic cell membranes $K^+$ cations.** Fluorescence intensity profiles of resorufin production when catalysed by membrane-bound hemin/G-quadruplex receptors localised in lipid domains of synthetic cell models for n = 3 repeats. Synthetic cell functionalisation was carried out in buffered solutions with added KCl and glucose (1×TE + 100 mM KCl + 87 mM glucose). Blue arrows indicate the addition of $H_2O_2$ to trigger peroxidation.

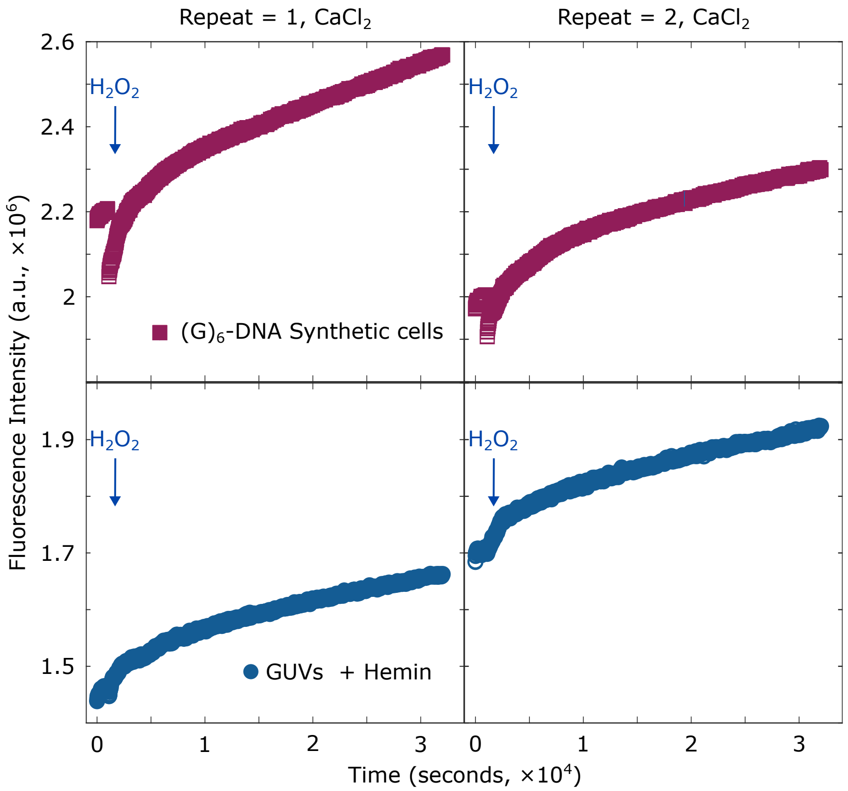


Figure S25: **Peroxidation activity localised to lipid domains of synthetic cell membranes $Ca^{+}$ cations.** Fluorescence intensity profiles of resorufin production when catalysed by membrane-bound hemin/G-quadruplex receptors localised in lipid domains of synthetic cell models for n = 2 repeats. Synthetic cell functionalisation was carried out in buffered solutions with added $CaCl_2$ and glucose (1×TE + 2 mM $CaCl_2$ + 281 mM glucose). Blue arrows indicate the addition of $H_2O_2$ to trigger peroxidation.

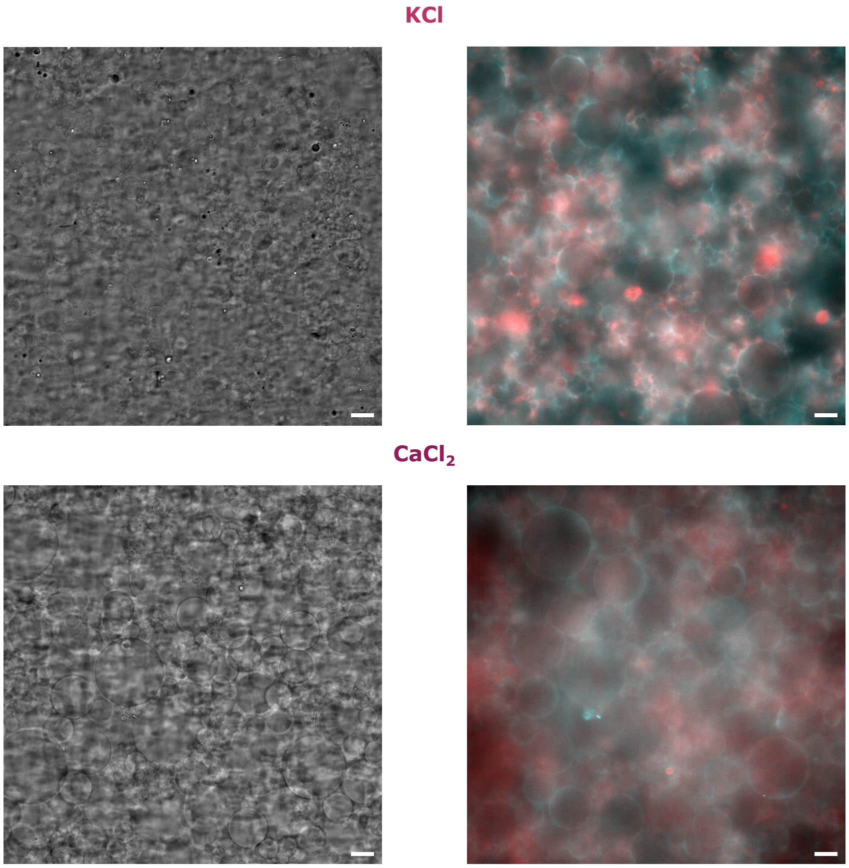


Figure S26: **Synthetic cells retain structural integrity after peroxidation.** Representative brightfield (left) and epi-fluorescence micrographs of DNA-functionalised synthetic cells in the presence of KCl (top) or $CaCl_2$ (bottom). Micrographs were acquired after the peroxidation reactions monitored with fluorimetry in Figs. 4e (main text), S16 and S17, showing their structural integrity following the conversion of AmplexRed to resorufin when triggered by $H_2O_2$. In the case of epifluorescence micrographs, TexasRed-DHPE signal, which stains $L_d$ domains, is shown in red, while the fluorescence signal of DNA nano-devices is shown in black. Scale bars = 20 $\mu$m.

### Supplementary Tables

Table S1: **DNA nano-devices and oligonucleotides that comprise them.** Each monomeric nanostructure is self-assembled through a quenching temperature ramp as discussed in the Experimental Section at a final concentration of 2 $\mu$M and then used for vesicle functionalisation. Constructs require strands in 1:1 stochiometric ratios.

| **Construct** | **Strands** |
|---|---|
| $Anchor_{dC}$ | $D_{bb,chol}$ + $D_{b,chol}$ + Spacer + (G)-rich strand (added after overnight incubation) |

Table S2: **Sequences of oligonucleotide strands.** Abbreviations: TEG: triethyleneglycol.

| **Strand** | **Sequence (5' → 3')** |
|---|---|
| $D_{b,chol}$ | CCAACACAACAACAAACCCGTTCCGACATAGAACCG/Cholesterol-TEG/ |
| $D_{bb,chol}$ | /Cholesteryl-TEG/CGGTTCTATGTCGGAACG |
| Spacer | GGTTTGTTGTTGTGTTGGAAACTGACAGACCTATTCGC |
| $(G)_6$-strand | TGGGGGGTTTTTTTTTGCGAATAGGTCTGTCAGTTT |
| $(G)_5$-strand | TGGGGGTTTTTTTTTGCGAATAGGTCTGTCAGTTT |
| $(G)_4$-strand | TGGGGTTTTTTTTTTGCGAATAGGTCTGTCAGTTT GATGAGAGAGC |
| $(T)_6$-strand | TTTTTTTTTTTTTTTTGCGAATAGGTCTGTCAGTTT |
| $(G)_6$-strand-Toehold | TGGGGGGTTTTTTTTTGCGAATAGGTCTGTCAGTTTGTTGTA |
| Spacer-Fluorophore | GGTTTGTTGTTGTGT/FAM-T/GGAAACTGACAGACCTATTCGC |
| Invader | TACAACAAACTGACAGACCTATTCGC |

Table S3: **Physiologically-relevant concentrations of cations.** Typical ranges of concentrations found in extracellular fluids and intracellular environments (e.g. cytosol, mithochondria, lysosome, endoplasmic reticulum, Golgi apparatus).[10]

| **Cation** | **Intracellular environment (mM)** | **Extracellular environment (mM)** |
|---|---|---|
| $K^+$ | $\sim 30 - 150$ | $\sim 3.5 - 5$ |
| $Na^+$ | $\sim 10 - 20$ | $\sim 135 - 140$ |
| $Ca^{2+}$ | $\sim 2$ | $< 1$ |
| $Mg^{2+}$ | $\sim 4 - 20$ | $0.5 - 1$ |

Table S4: **Statistical significance of fold-change in lipid domain partitioning.** Significance $p$-values obtained for each cationic condition using a non-parametric, one-tailed, Wilcoxon-signed test to assess the statistical significance of $K_\mathrm{p}^*$ values being higher than 1.

| **Cationic condition** | **$p$–value** |
|---|---|
| $K^+$ | $3.7 \times 10^{-162}$ |
| $Na^+$ | $1.6 \times 10^{-123}$ |
| $Li^+$ | $5.8 \times 10^{-39}$ |
| $Ca^{2+}$ | $1.6 \times 10^{-16}$ |
| $Mg^{2+}$ | $3.9 \times 10^{-47}$ |

Table S5: **Statistical correlation and significance between GUV size and DNA nanostructure partitioning** Spearman correlation coefficients ($\rho$) and $p-$values for GUV populations in each cationic composition before and after the addition of $(G)_6$-strands.

| **Cation** | **State** | **Spearman $\rho$** | **$p$-value** |
|---|---|---|---|
| $Na^+$ | Before | -0.24846 | 0.23108 |
| $K^+$ | Before | -0.43706 | 0.02004 |
| $Li^+$ | Before | 0.37013 | 0.04811 |
| $Mg^{2+}$ | Before | 0.34733 | 0.06002 |
| $Ca^{2+}$ | Before | 0.34341 | 0.04676 |
| $Na^+$ | After | 0.18595 | 0.32519 |
| $K^+$ | After | -0.17068 | 0.32695 |
| $Li^+$ | After | 0.18222 | 0.45528 |
| $Mg^{2+}$ | After | -0.24922 | 0.26336 |
| $Ca^{2+}$ | After | 0.55323 | 0.00152 |
